# Blooms like it hot, but mussels do not: Influence of invasive quagga mussels on cyanobacteria during summer

**DOI:** 10.64898/2026.02.21.707163

**Authors:** Jonas Mauch, Maider Erize Gardoki, Raphael Neiling, Jan Köhler, Jordan Facey, Sabine Hilt

**Author notes:** Email Addresses.

## Abstract

Quagga mussels (*Dreissena rostriformis bugensis*) are among the most impactful invaders in freshwaters of the Northern Hemisphere. As filter-feeders, they can reduce harmful algal blooms (HABs), but their effects are expected to be dependent on cyanobacteria species and water temperature. However, conclusive studies on these traits and their combination are lacking. Here, we combined laboratory experiments with an analysis of long-term data from a temperate shallow lake 10 years before and after quagga mussel invasion, respectively. We tested the hypotheses that quagga mussel filtration rates in the laboratory would 1) vary among common cyanobacteria species and 2) decrease above a critical temperature. Regarding the field data, we expected that 3) quagga mussels can reduce the summer biovolume of palatable cyanobacteria, but that 4) this effect disappears above a critical temperature. Our results support all four hypotheses. In laboratory experiments, *Dolichospermum flos-aquae* was classified as palatable to quagga mussels, while *Aphanizomenon flos-aquae, Anabaenopsis elenkinii* and *Microcystis aeruginosa* were less-palatable cyanobacteria. Filtration rates decreased above 28.9°C (CI: 27.6–30.2°C) with mussels dying at 32°C. Our long-term lake data show that cyanobacteria biovolumes were lower after quagga mussel invasion, but only below 27.7°C (CI: 26.9–28.4°C), confirming a critical thermal window for quagga mussel filtration. Global warming will therefore facilitate HABs by increasing the growth rates of cyanobacteria and reducing the filtration rates of quagga mussels above critical summer water temperatures, which are increasingly being reached in invaded lakes. This critical thermal window must be considered when making HAB predictions.

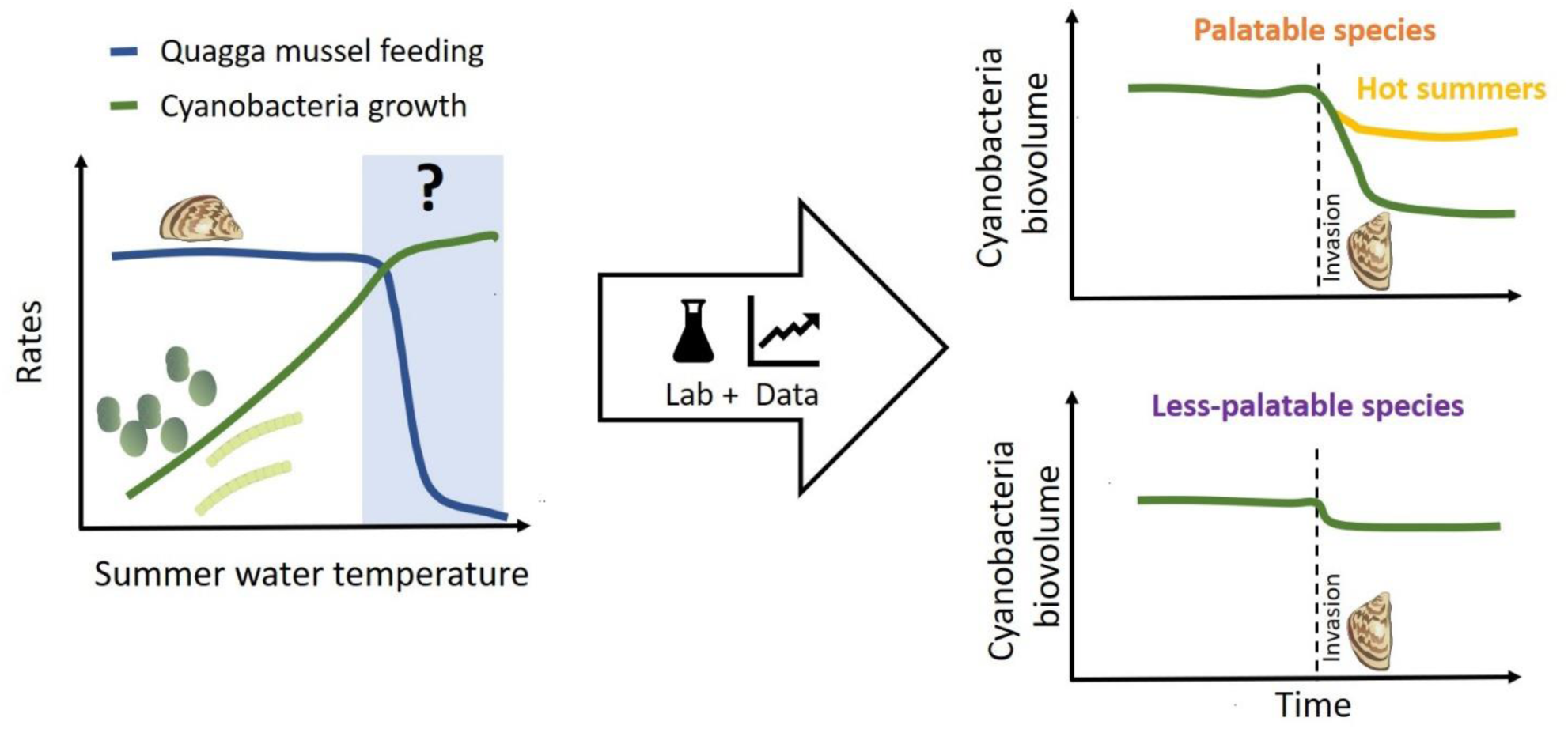

## 1. Introduction

Cyanobacteria are the most notorious producers of harmful algal blooms (HABs) in freshwaters (Paerl et al. 2001; Wang et al. 2021), a serious environmental issue due to their impact on ecosystems, as well as on animal and human health (Paerl and Otten 2013; Chorus and Welker 2021; Igwaran et al. 2024). Different typical harmful cyanobacteria species, such as *Microcystis aeruginosa, Aphanizomenon flos-aquae* and *Dolichospermum flos-aquae* (formerly classified as planktonic *Anabaena flos-aquae*), vary in their toxin production and morphology (Supplementary Table S2). They occur as single cells that can aggregate into colonies (*M. aeruginosa*), as single filaments (*D. flos-aquae*) or as filaments that aggregate into rafts (*A. flos-aquae*), which affects their resistance to grazing (Paerl et al. 2001; Wang et al. 2024). Although cyanobacteria blooms can also occur under relatively cold conditions (Reinl et al. 2023), most cyanobacteria generally grow faster than other phytoplankton groups at higher temperatures (above 25°C), making climate warming a powerful catalyst for cyanobacteria blooms (Paerl and Huisman 2008; O’Neil et al. 2012). Global mapping has revealed a significant increase in algal blooms (including cyanobacteria) in freshwater lakes over the last decade (Hou et al. 2022).

One of the major loss processes of HABs is filtration by dreissenid mussels (Harris et al. 2024). Two species of dreissenid mussels have been reported to be invasive: the quagga mussel (*Dreissena rostriformis bugensis*) and the zebra mussel (*Dreissena polymorpha*), whereas in recent years, quagga mussels have overtaken the previous zebra mussel invasion in many lakes (Nalepa 2010; Karatayev et al. 2015). Their invasion often results in increased water clarity and decreased chlorophyll (chl) *a* and phytoplankton biovolume in lakes (Higgins and Vander Zanden 2010; Karatayev and Burlakova 2022). While some studies have reported a strong decrease in total cyanobacteria biovolume after dreissenid mussel invasion (Nicholls et al. 2002; Kirsch and Dzialowski 2012; Reynolds and Aldridge 2021), others have demonstrated an increase in specific cyanobacteria species, including *Microcystis* sp. (Vanderploeg et al. 2001; Raikow et al. 2004; Knoll et al. 2008).

Dreissenid mussels can actively eject cyanobacteria cells from undesirable species (Vanderploeg et al. 2001; Tang et al. 2014). Several studies suggest that this selective feeding behaviour is influenced by the morphological characteristics of the cyanobacteria such as colony formation and the presence of gelatinous layers or thick cell walls (Vanderploeg et al. 2001; Dionisio Pires and Van Donk 2002; Dionisio Pires et al. 2005). However, toxicity does not appear to be a decisive factor (Vanderploeg et al. 2001; Dionisio Pires et al. 2005). Water temperature may also affect the impact of dreissenid mussels on cyanobacteria. While the filtration rates of zebra and quagga mussels remain relatively constant within a temperature range of approximately 10–24°C (Reeders and de Vaate 1990; Diggins 2001; Xia et al. 2021), a decrease in filtration rates has been observed in zebra mussels at temperatures above 24°C (Aldridge et al. 1995; Lei et al. 1996). There is only one study available for quagga mussel filtration rates above 24°C (Gopalakrishnan and Kashian 2020), which does not clearly indicate a critical upper temperature limit for filtration. The average summer (July 15–August 31) epilimnetic temperatures in 345 northern hemisphere lakes (22.0 ± 3.0°C) already reach or exceed this value (Zhou et al. 2023), and this will become more common in the near future due to warming and increased heatwave frequency (Woolway et al. 2021). For the Great Lakes, water temperatures are projected to increase by 3.3–6.0°C by the end of the century (Zhang et al. 2020). Similarly, European temperate lakes are warming at a rate of 0.1-0.3°C per decade, with the steepest warming trends observed in shallower lakes (Shatwell et al. 2019; Schwefel et al. 2025). Climate warming is projected to particularly increase the intensity and absolute magnitude of heatwaves and peak water temperatures in lakes (Woolway et al. 2020, 2021), which may push dreissenid mussels close to their upper thermal limits (Aldridge et al. 1995; Spidle et al. 1995). Heatwaves in freshwater systems can strongly affect biological processes across trophic levels, often producing more acute effects than moderate continuous warming (Polazzo et al. 2022; Zhang et al. 2022). Many lakes in the northern hemisphere are already experiencing, or are vulnerable to, both cyanobacteria blooms (Visser et al. 2016) and quagga mussel invasions (Quinn et al. 2014; Haltiner et al. 2022; Kraemer et al. 2023) and climate change is projected to increase the extent of quagga mussel invasions, potentially affecting thousands additional lakes (Gallardo and Aldridge 2025). The effect of rising summer water temperatures on quagga mussels’ filtration on cyanobacteria is not well understood, making it difficult to predict future cyanobacteria blooms in these lakes.

Here, we conducted filtration rate experiments with quagga mussels on four species of common bloom-forming cyanobacteria with different morphologies, two species of green algae and mixed cyanobacteria communities from a lake, at temperatures ranging from 24 to 33°C. We also analysed long-term data from a temperate shallow lake. We compared the summer biovolumes of abundant cyanobacteria species along a temperature gradient over 10 years before and after the invasion of quagga mussels, respectively. We hypothesized that quagga mussel filtration rates in laboratory experiments would 1) vary among different common bloom-forming cyanobacteria species and 2) significantly decrease above a critical temperature. Regarding the field data, we expected that 3) quagga mussels can reduce the summer biovolume of palatable cyanobacteria, but that 4) this effect disappears above a critical temperature.

## 2. Materials and methods

### 2.1 Filtration rate experiments

We determined the filtration rates of quagga mussels at various temperatures ranging from 24 to 33°C using: 1) single-strain cultures of the four most abundant cyanobacteria species in eutrophic Lake Müggelsee, which was used for long-term data analyses (further information below), 2) two green algae species known to be palatable to dreissenid mussels (Ackerman 1999; Vanderploeg et al. 2013) and 3) mixed lake samples collected during late summer cyanobacteria blooms in Lake Müggelsee.

#### 2.1.1. Phytoplankton

The phytoplankton species *Dolichospermum flos-aquae*, *Anabaenopsis elenkinii*, *Chlorella vulgaris* and *Microcystis aeruginosa* were obtained from different culture collections (Supplementary Table S1). Cultures of the green algae *Acutodesmus obliquus* were originally obtained from SAG, but have been cultivated in our laboratory since 2014. The cyanobacterium *Aphanizomenon flos-aquae* was isolated from Lake Müggelsee in the summer 2023. We used species-specific growth media (Supplementary Table S1) as provided by the culture collections for optimal cultivation. To address the potential impact on mussel filtration, we included media type as an additional fixed effect in a general linear model (GLM), which revealed that media had no significant effect on filtration rates (F = 0.25, p = 0.615; see Supplementary Table S8). The effect of the medium type was therefore excluded from further GLMs. For the sake of simplicity, all phytoplankton species are further referred to by their genus names (*Dolichospermum, Anabaenopsis, Aphanizomenon, Microcystis, Acutodesmus, Chlorella*). The four cyanobacteria strains tested are all known from literature (Supplementary Tables S1 and S2) to potentially produce toxins, but differ in their morphology, colony formation and presence of gelatinous layers (Supplementary Table S2). Previous studies have demonstrated that dreissenid mussels filter toxic and non-toxic cyanobacteria strains at similar rates, indicating that toxin production is not the primary determinant of filtration selectivity (Dionisio Pires et al. 2005; Vanderploeg et al. 2001). All phytoplankton strains were grown in batch cultures at a constant temperature of 23°C, a light intensity of approximately 100 µE m^-2^ s^-1^ and a 16:8 h day:night cycle, in a Binder KBW 720 incubator (Germany). Batches were cultivated in Erlenmeyer flasks, increasing in size from 50 mL to 5 L as necessary, and agitated at 90 rpm on GFL 3005 orbital shakers (Gesellschaft für Labortechnik mbH, Germany). We added phytoplankton-specific growth media on a weekly basis or more frequently if necessary, to maintain exponential growth phase. The cultivation period for all phytoplankton species varied depending on the growth rate of the different cultures and ranged from several days to several weeks until sufficient biomass was reached.

As cyanobacteria colony formation can differ between laboratory cultures and *in situ* lake conditions, we also measured quagga mussel filtration rates on cyanobacteria collected during summer blooms in Lake Müggelsee dominated by *Microcystis aeruginosa* and *Aphanizomenon flos-aquae.* Two different cyanobacteria concentrations were tested, ‘Lake Müggelsee high’ (30 µg L^-1^ chl *a*) and ‘Lake Müggelsee low’ (15 µg L^-1^ chl *a*). These were sampled by filtering lake water through a 30 μm plankton net (Apstein 50, Hydro-Bios, Germany) from a boat. In the laboratory, we filtered the entire sample (approximately 100 L) through a 200 μm plankton net to remove large zooplankton. Samples were collected one day prior to the filtration experiment and stored in a large plastic container at room temperature (approximately 20°C). Immediately prior to the experiment, we measured chl *a* concentrations in the storage tank using a Phyto-PAM fluorometer (Heinz Walz GmbH, Effeltrich, Germany) and identified phytoplankton taxa via microscopic counting to determine the bloom composition (for detailed results see Supplementary Tables S3 and S4). In addition, chl *a* was measured by high-performance liquid chromatography (HPLC). The HPLC system (Waters Alliance) consisted of a WATERS Separation Module 2695, a WATERS Photodiode Array Detector 2996 and a Symmetry Shield RP18 column (100 Å, 3.5 µm, 4.6 mm × 150 mm, 1/pk). The injected volume was 60 µL sample and 30 µL ultrapure water. This method is described in detail by Shatwell et al. (2012).

#### 2.1.2. Quagga mussel sampling and storage

Quagga mussels were collected by hand from the north-western shore of Lake Müggelsee (52° 26’ 46.0” N, 13° 37’ 57.5” E), at a depth of 1.5 m and transported to the laboratory within two hours. Mussels with a shell length of approximately 15-30 mm were selected to exclude younger individuals that had not previously been exposed to summer cyanobacteria blooms. Approximately 300-350 individuals were kept at room temperature (∼ 20°C) in plastic containers (height: 30 cm, width: 54 cm, length: 78 cm) with 50 L of tap water (bank filtrated water from Lake Müggelsee, non-chlorinated and copper-free) aerated with air pumps (AquaForte Air Pump HI-Flow V-20, Germany) and air diffusers. The mussels were collected from the lake at least three weeks before the start of each experiment (Supplementary Table S1) and gradually adapted to tap water, temperatures and food. Although seasonal differences in quagga mussel filtration rates have been reported by Diggins (2001), these were attributed to differences in water temperature (8-22°C). We allowed at least 3 weeks for the mussels to adapt to laboratory conditions, so we assume that the timing of the mussel sampling did not influence their filtration rates. Initially, the mussels were kept in a mixture of 50% lake water and 50% tap water for one week, then to 25% lake water and 75% tap water, and finally to 100% tap water for one more week. The mussels were fed for three hours weekly with one teaspoon (0.55 g ± 0.1 g) of crushed TetraMin flakes (Tetra GmbH, Germany), and two teaspoons (3.8 g ± 0.25 g) of dried *Spirulina* algae (Naturwaren-Niederrhein GmbH, Germany), according to a long-established method developed by colleagues at the University of Duisburg-Essen. The water was then changed after feeding. Dead mussels and empty shells were removed within 2 days after collection from the lake and again after each feeding.

#### 2.1.3. Experimental set-up

The filtration rates of quagga mussels were tested at 24–33°C in 1°C increments for *Acutodesmus* as a model organism and at 24–32°C in 2°C increments for all other species (due to time and space constraints) according to the method described by Vanderploeg et al. (2001). Temperatures were randomly assigned to five incubators (3x Binder KBW series, Germany and 2x Pol-Eko Aparatura, Smart Pro, Poland). The mussels were fed with half a teaspoon (0.27 ± 0.05 g) of crushed TetraMin flakes, and one teaspoon (1.9 ± 0.13 g) of *Spirulina* algae (50% of the amount fed during acclimation) 17 h prior to the start of each experiment. One day before an experiment, we filled glass vessels (height: 29 cm, diameter: 9.5 cm) with 2 L of tap water and placed them into the incubators to reach the appropriate temperature. We randomly selected four quagga mussels (length: 13-27 mm, Supplementary Figure S1) per vessel and acclimated them to the glass vessels 1 h before the start of the filtration rate experiments. To start the experiments, we used Eppendorf pipettes to add the algae/cyanobacteria with an initial concentration of about 20 µg chl *a* L^-1^, as determined by the Phyto-PAM fluorometer (Heinz Walz GmbH, Effeltrich, Germany). We collected 10 mL water samples from all vessels at the beginning and at the end of the experiments after 3 h and measured them with PhytoPAM or HPLC (only for Lake Müggelsee experiments). Four treatments, each containing mussels, phytoplankton and tap water, and four controls, containing only phytoplankton and tap water were used at each temperature. All experiments were performed in the absence of light in the incubators, to prevent phytoplankton growth. Mussel filtration was ensured by checking for siphon activity 10 min after the start of the experiment.

Mixed lake phytoplankton samples were treated similarly, but the vessels were not filled a day before to reach their temperatures (as in the single species algae/cyanobacteria experiments). The vessels were filled with lake samples 2-3 h prior to the experiment to reach the target temperature, in the incubators and started the experiment by adding the mussels, which then were exposed to the samples for 3 h. All experiments for the different temperatures in the incubators were started at 15 min intervals to allow sufficient time for sampling in between.

Based on the chl *a* concentrations at the beginning and at the end of the experiments, we calculated filtration (clearance) rates of quagga mussels using the following equation from Vanderploeg et al. (2001):

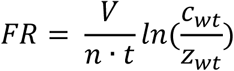

Where: FR = Filtration rate (mL mg^-1^ h^-1^); V = total water volume in vessel (mL); n = mussel ash-free dry weight (AFDW) in vessel (mg); t = duration of experiment (h); c_wt_ = mean concentration of algae in control vessels at the end of the experiment (μg L^-1^); and z_wt_ = concentration of algae in a mussel vessel at the end of the experiment (μg L^-1^).

After the experiment, the mussel shell lengths were measured with a digital caliper (precision = 0.5 mm) and stored in a freezer at –20°C. To normalize the filtration rates, we determined the shell-free dry weight (SFDW) and ash-free dry weight (AFDW) of the mussels according to Nalepa et al. (2010). For the measurement of SFDW, the soft tissue was separated from the shells while still frozen, placed in pre-weighed aluminium cups, dried in an oven (Heraeus Instruments T 6030, Germany) at 60°C for 48 h and weighed. To determine AFDW, dried tissue was combusted at 450°C for 1 h (Nabertherm N 11 oven, Germany) and weighed to an accuracy of 0.01 mg. A few individual samples were destroyed in the process, so we used the length to dry weight ratio of all the measurements in Supplementary Figure S1 to calculate AFDW from shell length:

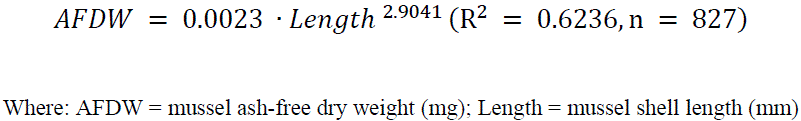

#### 2.1.4. Statistical analysis of filtration rate experiments

To evaluate species-specific differences in filtration rates and the effects of temperature, we employed a GLM with a Gaussian error distribution. The model included cyanobacteria genus (four levels: *Dolichospermum*, *Anabaenopsis*, *Aphanizomenon*, *Microcystis*), temperature (five levels: 24, 26, 28, 30, 32°C), and their interaction as fixed effects, to test for genus-temperature interactions. We used Type III ANOVA (likelihood ratio tests) to quantify main and interaction effects. Before fitting the GLM, we visually assessed normality of residuals using quantile-quantile (qq)-plots. Homogeneity of variance was assessed using Levene’s test and by inspecting residual plots against the fitted values. We evaluated the model fit using McFadden’s pseudo-R² and Akaike Information Criterion (AIC).

To identify specific pairwise differences between cyanobacteria genera at each temperature, we conducted Tukey-adjusted post hoc contrasts using estimated marginal means (’emmeans’ package, Lenth et al. 2025). These contrasts yielded p-values and standardised effect sizes (Cohen’s d) for all pairwise species comparisons at each temperature. Following convention, we interpreted the effect sizes as negligible (d < 0.2), small (d = 0.2–0.5), medium (d = 0.5–0.8) or large (d > 0.8).

To determine the critical temperature at which the mussel filtration rates significantly declined for palatable species, we fitted a segmented regression model using the segmented package (version 1.3-4, Muggeo 2003). This approach allowed us to identify the breakpoint (BP) temperature where the relationship between temperature and filtration rate changed from one linear segment to another. The model was applied to the combined filtration rate data of palatable cyanobacteria and green algae (*Dolichospermum*, *Chlorella*, and *Acutodesmus*), with temperature as the continuous predictor. The segmented regression algorithm iteratively estimated the BP location by minimising the residual sum of squares. We reported the estimated BP temperature alongside its 95% confidence interval.

### 2.2. Long-term data analysis

#### 2.2.1. Study site and water sampling

Lake Müggelsee (52° 26’ 0” N, 13° 39’ 0” E) is a shallow (mean depth 4.9 m, maximum depth 8 m), polymictic lake in Berlin (Germany). It covers a surface area of 7.3 km^2^, has a volume of 36,560,000 m^3^ and the River Spree flows through the lake with a theoretical retention time of 6–12 weeks. Its catchment area of 7,000 km^2^ consists of 36% forest and 42% agricultural land, populated by 720,000 people. In this lake, the primary growth-limiting nutrient for phytoplankton in the spring is phosphorus (P), while nitrogen (N) becomes limiting during the summer months due to high P release from the sediment, which is common in shallow polymictic lakes (Shatwell and Köhler 2019). Following a reduction in nutrient loading during the 1990s, aquatic macrophytes have become increasingly established in the lake (Hilt et al. 2013). The earliest report of invasive zebra mussels in Lake Müggelsee dates back to the early 20th century (Arndt et al. 1993), but their coverage remained low, because most of the lake is covered in soft sediment that zebra mussels cannot colonise. During a monitoring of 8 transects around the lake by scuba divers in 2011, dreissenid mussels comprised only *D. polymorpha* and 50% of the investigated 24 locations (three depth zones (0-1 m, 1-2 m, 2-4 m) at each of the eight transects) did not have any dreissenid mussels. Quagga mussels invaded the lake around 2012 (Wegner et al. 2019). In 2017, the abundance of dreissenid mussels in the lake had increased about 10fold compared to 2011 and quagga mussels had reached a proportion of 97.3% (zebra mussels 2.7%), covering about one third of the total lake area with a maximum density of almost 46,000 mussels m^-^² (Wegner et al. 2019). As part of a long-term monitoring program (IGB 2025), volumetrically weighted integrated water samples (∼105 L in total) from 21 subsamples from five different locations on the lake (Driescher et al. 1993; Shatwell and Köhler 2019) were collected weekly during the summer using a Friedinger sampler for analysis of nutrients and phytoplankton.

#### 2.2.2. Nutrient concentrations

Phosphorus analysis is based on Murphy and Riley (1962) according to EN ISO 6878. Unfiltered samples were used for the determination of total phosphorus (TP). For each sample, 25 mL were acidified with 2 mL of 10N H_2_SO_4_ solution. 1 mL H_2_O_2_ (20 %) was added and then heated in a digestion heating block at 150°C for at least 10 hours. After adjusting the pH to ∼ 7 with sodium hydroxide solution, 2 mL of molybdate sulfuric acid and 0.5 mL of 10 % ascorbic acid were added and the molybdenum blue was measured photometrically using a Cary 60 UV-VIS spectrophotometer in a 5 cm quartz cuvette. Soluble reactive phosphorus (SRP) was determined using a Continuous Segmented Flow Analyzer (FSA) in the Auto Analyzer from SEAL Analytical GmbH. The same colorimetric method as for TP was used. Samples were pre-filtered at 0.45 μm and 12 mL sample was acidified with 150 μL 2M HCl for preservation purposes.

Total nitrogen (TN) concentrations were measured after catalytic oxidation as chemoluminescence in a Shimadzu TOC/TN analyser according to DIN EN 1484. To ensure a pH < 2, 100µl of 2M HCl was added per 10mL sample. Dissolved inorganic nitrogen (DIN) concentrations were calculated as the sum of nitrate (NO_3_-N), nitrite (NO_2_-N) and ammonium (NH_4_-N) concentrations. For this purpose, the samples were pre-filtered with 0.45 μm pore size and 12 mL of the sample was preserved with 150 μL 2 M HCl. NH_4_ was determined by colorimetric measurement via FSA according to DIN 38 406. Sodium salicylate and dichloroisocyanuric acid reacted to form a blue color complex, which was measured photometrically. NO_3_ was determined according to ISO 13395 / DIN 38 405 in the same device using a colorimetric measurement with sulphanilamide. NO_3_ was reduced with cadmium segment to nitrite and after reaction with sulphanilamide the resulting azo dye was determined photometrically. NO_2_ was measured according to the steps in NO_3_.

#### 2.2.3. Cyanobacteria

Samples of 50 mL of lake water were fixed with Lugol’s solution in preparation for cyanobacteria analysis under an inverted microscope. The abundances of individual species or algal groups were counted and the corresponding dimensions were measured. Biovolumes were calculated by assuming the organisms were simple geometric shapes. Twenty individuals of the most abundant taxa were measured, with more than 20 individuals counted and all individuals of the remaining taxa were measured.

#### 2.2.4. Water temperature

Water temperatures in Lake Müggelsee are recorded at a monitoring station (52° 26’ 46.3” N, 13° 38’ 60.0” E; total depth at this location is 5 m). From 2002-2003, the temperature was measured using a thermistor chain (Energie-Anlagen-Berlin, semiconductor sensors AD592) at 5-min intervals at a depth of 0.5 m. From 2003-2021, the water temperature was measured with a multi-parameter probe (YSI6600, YSI Incorporated, USA) at a water depth of 0.5 m.

To compare the water temperatures measured at the monitoring station in the pelagic with those of littoral areas, additional measurements were taken in the littoral zone, at a depth of 0.5 m (total water depth of approximately 1 m) at the north shore of the lake (52° 26’ 52.8” N, 13° 38’ 55.7” E) from July 01, 2023 until August 25, 2023 using two HOBO Temperature Loggers UA-002-64 (Onset, USA). We calculated the mean values from both HOBO loggers for our analysis.

#### 2.2.5. Data analysis and statistics

To compare field data from the decade before (2002-2011) and after (2012-2021) the quagga mussel invasion, we selected all long-term data from the three summer months (July-September). During this period, cyanobacteria blooms were frequently recorded, and phosphorus limitation could be ruled out in Lake Müggelsee (Shatwell and Köhler 2019). We tested all the data for normal distribution (via qq-plot) in order to select the suitable statistical tests.

The nutrient concentration data were log-transformed to improve normality and homoscedasticity prior to analysis. Levene’s tests were used to assess the homogeneity of variances for each nutrient in relation to invasion status (pre-quagga vs. post-quagga). To evaluate the overall differences in the multivariate nutrient profile between the invasion periods, a multivariate analysis of variance (MANOVA) was conducted, employing Pillai’s trace as the test statistic, given its resilience to deviations from multivariate normality and homogeneity of variance-covariance matrices. This included four dependent variables: soluble reactive phosphorus (SRP), total phosphorus (TP), dissolved inorganic nitrogen (DIN), and total nitrogen (TN), with invasion status as the independent variable. Following the MANOVA, univariate analyses of variance (ANOVAs) were conducted for each nutrient to identify those exhibiting significant changes between the pre– and post-invasion periods. To investigate relationships among nutrients and potential shifts following invasion, Pearson correlation coefficients were calculated for all nutrient pairs within the pre-invasion, post-invasion, and overall study periods, and visualized using correlograms to facilitate the interpretation of changing nutrient dynamics.

We assigned the maximum water temperature (based on hourly means) from the seven days prior to the weekly phytoplankton sampling to each cyanobacteria biovolume, except in Figure 5, where we selected the daily maximum water temperature (based on hourly means). In our study, we expected thermal maxima to more accurately capture the physiological stress that impairs the filtration capacity of mussels than mean temperatures.

For all long-term data analyses of cyanobacteria biovolume we excluded days on which total cyanobacteria biovolume was below 1 mm^3^ L^-1^ (except for Figure 5) in order to eliminate days on which cyanobacteria abundance was below the low-risk threshold, defined by the WHO (Lyche Solheim et al. 2023). All cyanobacteria species were grouped by genus. Biovolumes for each cyanobacteria genus were analysed using a generalized linear mixed model (GLMM) with zero-inflation, implemented with the ‘glmmTMB’ package (Brooks et al. 2017). The response variable, biovolume, was modelled using a gamma error distribution and a log-link function to accommodate its strictly positive nature, while the zero-inflation component accounted for any excess zeros. The model included invasion status (pre– and post-quagga) and cyanobacteria genus (*Dolichospermum*, *Anabaenopsis*, *Microcystis*, and *Aphanizomenon*) as fixed effects, along with their interaction term, in order to assess differential genus responses to invasion. A random intercept for sampling date was included to account for temporal clustering of observations collected on the same date. Model coefficients were interpreted on the log scale, with exponentiated estimates representing the multiplicative effect on biovolume. All diagnostic procedures and graphical assessments were conducted to ensure that the model met the necessary assumptions.

To assess the differential impact of quagga mussel invasion on cyanobacteria based on palatability, we categorised the four cyanobacteria genera into two functional groups: palatable (*Dolichospermum*) and less-palatable (*Aphanizomenon*, *Anabaenopsis*, and *Microcystis*). The categorization was based on filtration rate experiments (Figure 1). We then tested the influence of invasion on these palatable and less-palatable cyanobacteria groups using the identical GLMM method. Here, cyanobacteria biovolume was modelled as a function of invasion status and palatability group and their interaction, with a random intercept for sampling date.

**Figure 1:**
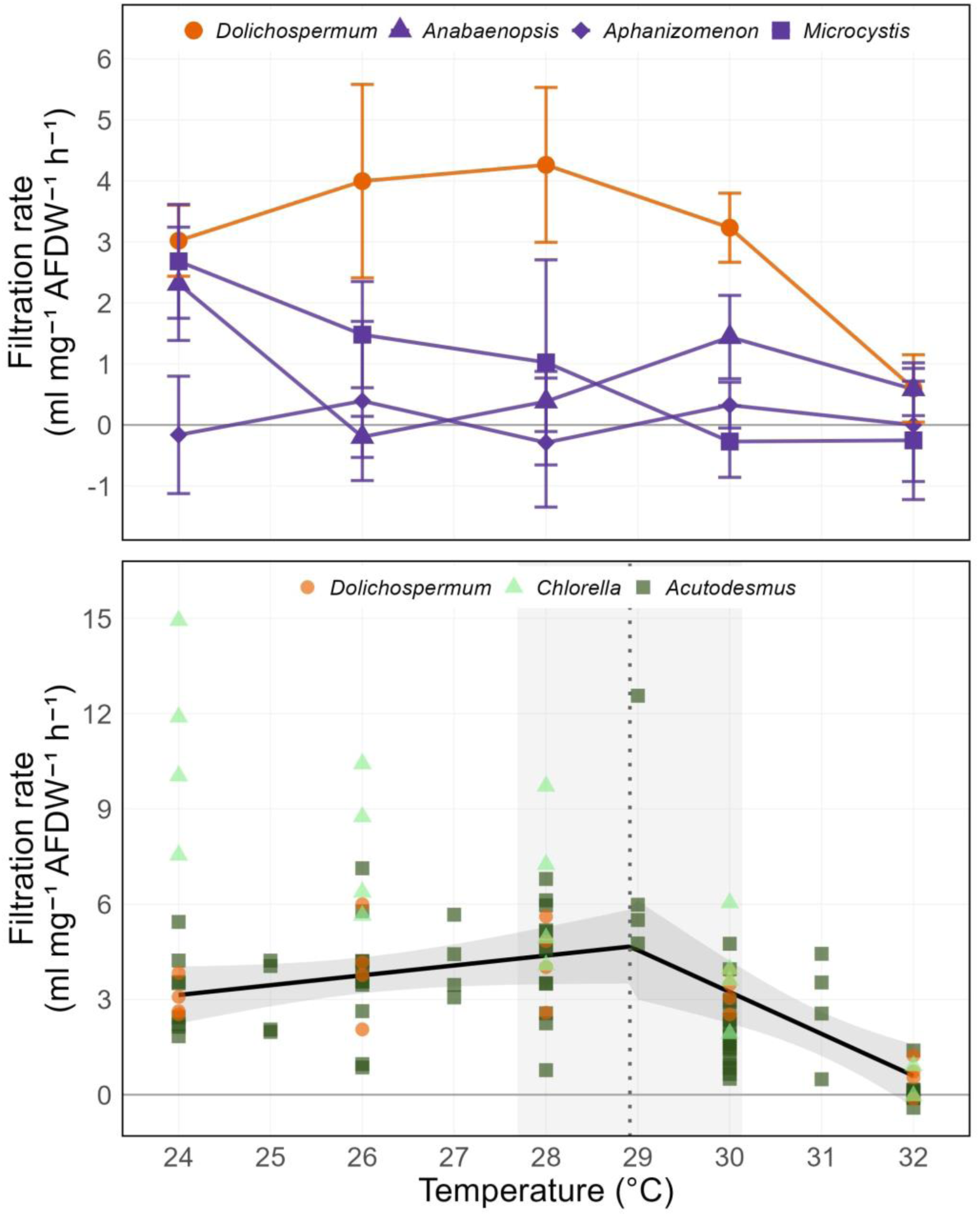
Filtration rates of quagga mussels on different phytoplankton species at various water temperatures (24–32 °C). The mean filtration rates (± standard deviation) are shown. (A) for palatable (orange: Dolichospermum flos-aquae) and less-palatable cyanobacteria (purple: Anabaenopsis elenkinii, Aphanizomenon flos-aquae, and Microcystis aeruginosa). Differences in filtration rates among cyanobacteria based on generalized linear model (GLM) analysis. Segmented regression analysis (B) of the filtration rates of quagga mussels on one palatable cyanobacteria species (orange: Dolichospermum flos-aquae) and two palatable green algae species (light/dark green: Chlorella vulgaris and Acutodesmus obliquus) with a breakpoint (black dotted line) at 28.9 °C (95% confidence interval: 27.7–30.1 °C, grey box). The black solid line represents the segmented regression model. For additional information see Supplementary Figure S2 and Tables S5-S7.

Segmented regression analyses and the Davies test were used to identify potential BPs and test for significant differences in the slope parameters for palatable and less-palatable cyanobacteria biovolume in relation to maximum water temperature before and after quagga mussel invasion. These statistical analyses were performed using the ‘segmented’ package (version 1.3-4; Muggeo 2003). The Davies test was employed specifically to assess the statistical significance of the identified break-points, providing a quantitative assessment of whether the slope change at the BP was statistically significant. To validate the assumptions of our linear regression model, we tested for homoscedasticity in the relationship between water temperature and cyanobacteria biovolume for palatable and less-palatable cyanobacteria before and after the invasion. A linear model was fitted to the data, and the residuals were analysed using both visual inspection and formal statistical testing. We performed the Breusch-Pagan test (Breusch and Pagan 1979) using the ‘lmtest’ package (Hothorn et al. 2022) to quantitatively assess whether the variance of the residuals remained constant across the range of predicted values.

We performed all statistical analyses using R statistical software version 4.3.0 (R Core Team 2023), and plotted figures using the ‘ggplot2’ (Wickham 2016), ‘cowplot’ (Wilke 2020) and ‘ggsignif’ (Ahlmann-Eltze and Patil 2021) packages. We chose significance levels as follows: n.s. (not significant): *p* ≥ 0.05; *: *p* < 0.05; **: *p* < 0.01; ***: *p* < 0.001.

## 3. Results

### 3.1. Filtration rate experiments

At temperatures of 32°C and 33°C, all quagga mussels were dead after 4 h (1 h acclimation followed by 3 h of experiment). Analyses of the effects of species and temperature on filtration rates (GLM with Type III ANOVA on four cyanobacteria species) revealed highly significant main effects for cyanobacteria species (χ² = 27.64, p < 0.001, η² = 0.11) and temperature (χ² = 37.03, p < 0.001, η² = 0.14). The strongest effect was the species × temperature interaction (χ² = 55.09, p < 0.001, η² = 0.20), indicating that species-specific filtration responses varied across temperature levels (see Supplementary Tables S6 and S7). At 24°C, filtration rates ranged from approximately 0–3.0 mL mg⁻¹ AFDW⁻¹ h⁻¹ across species (Figure 1A). While *Aphanizomenon* showed numerically lower rates than the other three species, these differences were not statistically significant (all pairwise contrasts p > 0.70). At 26-28°C, *Dolichospermum* exhibited higher filtration rates (∼4.0 mL mg⁻¹ AFDW⁻¹ h⁻¹) than *Anabaenopsis, Microcystis*, and *Aphanizomenon* (all < 1.0 mL mg⁻¹ AFDW⁻¹ h⁻¹; Cohen’s d = 2.5-4.2, all p < 0.01). Based on these differences, we categorised *Dolichospermum* as a palatable species and *Microcystis*, *Aphanizomenon*, and *Anabaenopsis* as less-palatable species for quagga mussels.

Both *Chlorella* and *Acutodesmus* exhibited high filtration rates within the temperature range of 24-28°C (4-15 mL mg⁻¹ AFDW⁻¹ h⁻¹; Figure 1A, B). Mixed lake cyanobacteria, dominated by *Microcystis* and *Aphanizomenon*, were not, or only negligibly, filtered by quagga mussels across all temperatures (Supplementary Figure S2G, H), confirming them as less-palatable species.

Segmented regression analysis of filtration rates of quagga mussels on three palatable phytoplankton species (*Dolichospermum*, *Chlorella*, *Acutodesmus*) at different temperatures revealed a decline in filtration rates within the range from 27.7 to 30.1°C (95% CI) with a BP at 28.9°C (shaded area and dotted line in Figure 1B). Below this temperature range, filtration rates remained relatively stable. Consistent with the complete mussel mortality observed at 32°C, all cyanobacteria species exhibited filtration rates near zero (0.25–0.60 mL mg AFDW⁻¹ h⁻¹), with no differences between species (all p > 0.59, Cohen’s d < 0.2).

### 3.2. Long-term data analysis

In Lake Müggelsee, a multivariate analysis of nutrient concentrations revealed a highly significant overall difference in nutrient composition between pre– and post-invasion periods (MANOVA: Pillai’s λ = 0.511, F = 60.12, p < 0.001). Follow-up univariate tests showed that DIN concentrations did not differ significantly between periods (F = 0.27, p = 0.60, η² = 0.001). However, TN (F = 61.52, p < 0.001, η² = 0.21), SRP (F = 63.25, p < 0.001, η² = 0.21), and TP (F = 156.23, p < 0.001, η² = 0.40) exhibited significantly lower concentrations in the post-invasion period (Figure 2, Supplementary Table S9 and S10).

**Figure 2:**
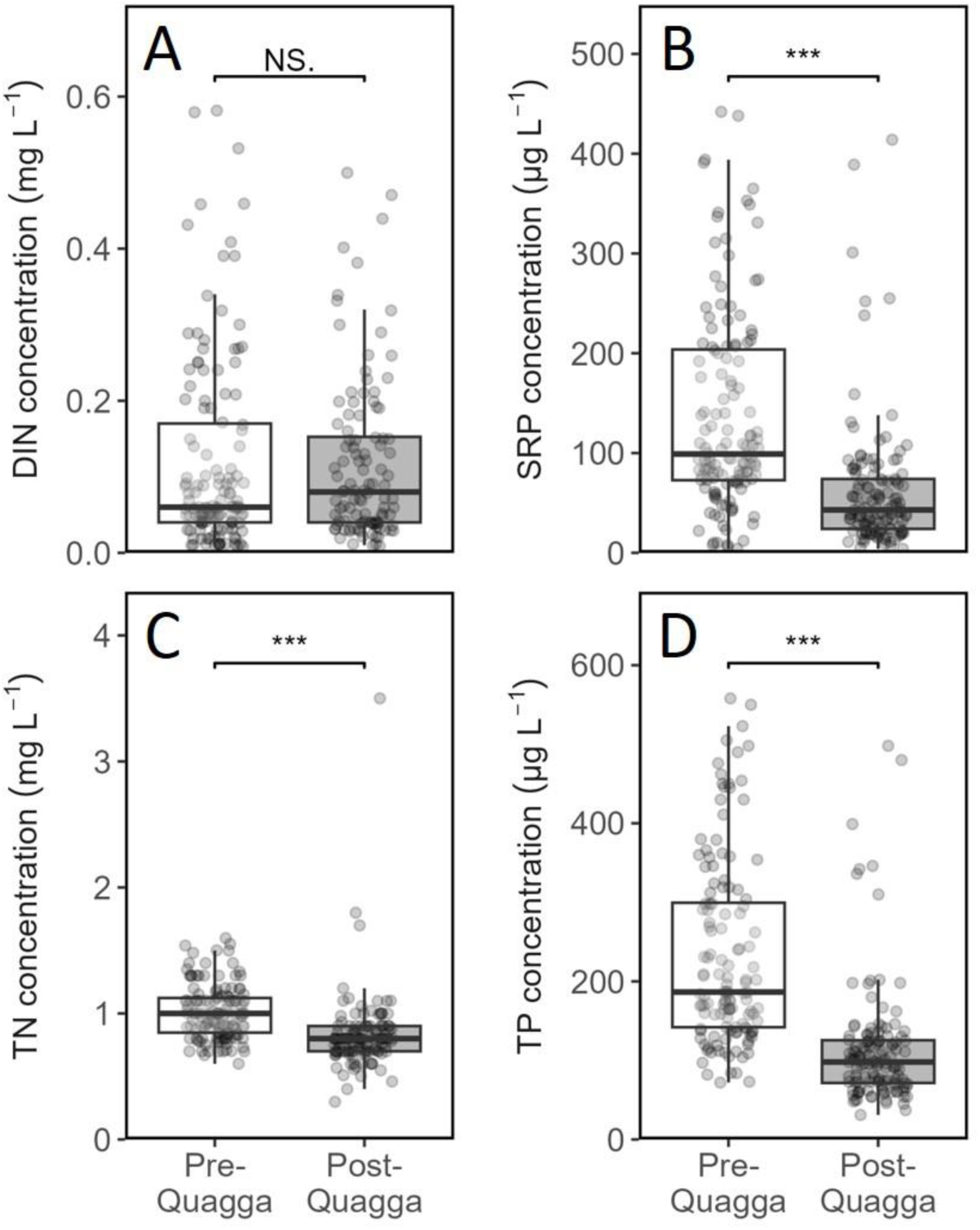
Summer (July-September) nutrient concentrations in Lake Müggelsee before (2002-2011, white) and after (2012-2021, grey) quagga mussel invasion. The concentrations are shown for: A: dissolved inorganic nitrogen (DIN), B: soluble reactive phosphorus (SRP), C: total nitrogen (TN), D: total phosphorus (TP). Comparisons between the two decades were performed using MANOVA (***: *p* < 0.001). For details see Supplementary Figure S3, S4 and Tables S9 and S10.

Cyanobacteria biovolumes in Lake Müggelsee during summer (Figure 3) were dominated by *Dolichospermum* both before and after quagga mussel invasion. The biovolumes of *Aphanizomenon* and *Microcystis* ranked second and third, while *Anabaenopsis* had the lowest biovolumes. After quagga mussel invasion, *Dolichospermum* exhibited a significantly lower biovolume (zero-inflated GLMM, p < 0.001), while the biovolumes of the other cyanobacteria did not change after the invasion (for details see Supplementary Figure S5 and Tables S11 and S12).

**Figure 3:**
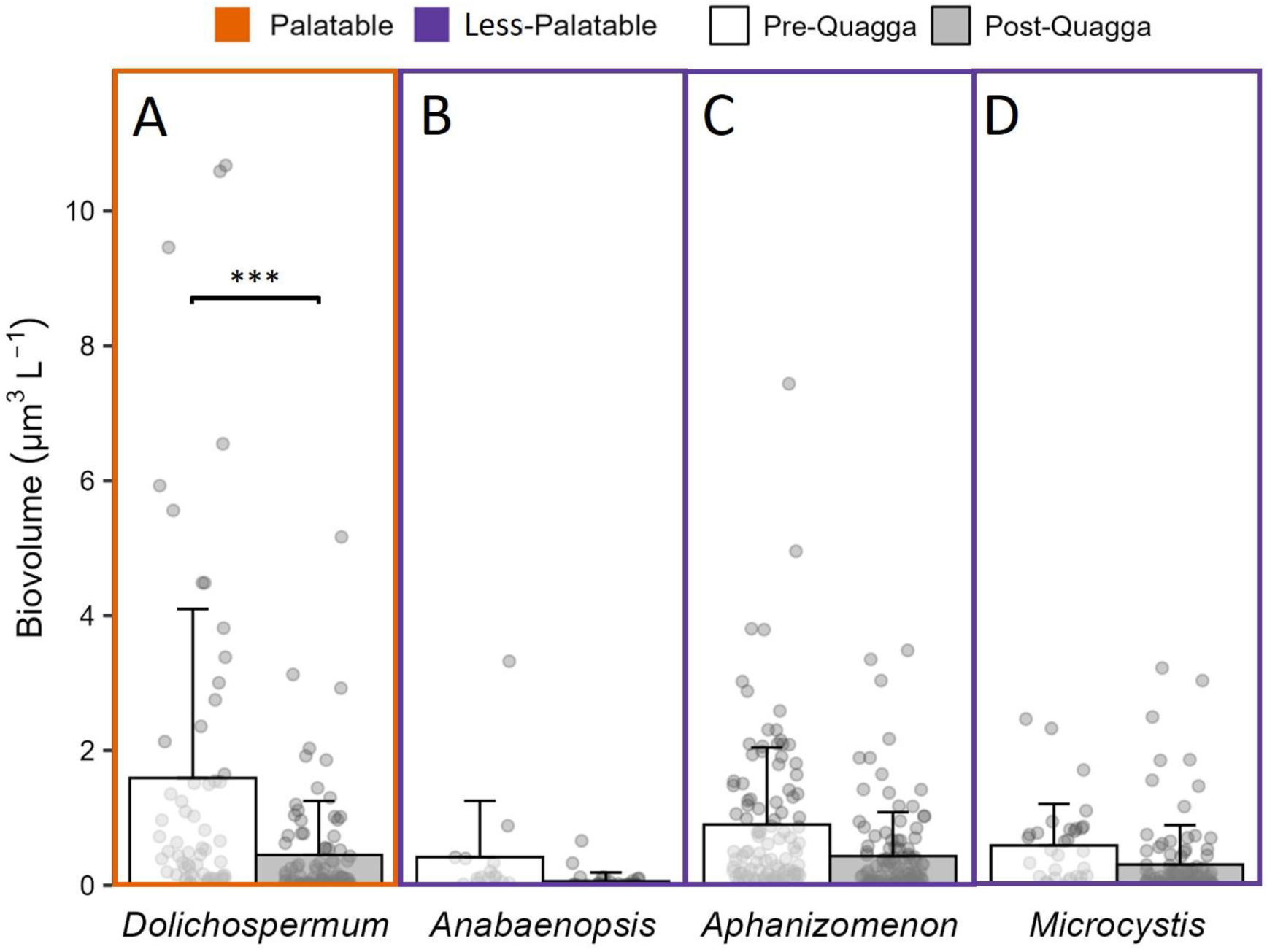
Summer (July-September) cyanobacteria biovolume of the four most abundant cyanobacteria species. (A: *Dolichospermum flos-aquae*, B: *Anabaenopsis elenkinii*, C: *Aphanizomenon flos-aquae* and D: *Microcystis aeruginosa*) in Lake Müggelsee before (Pre-Quagga: 2002-2011; white) and after (Post-Quagga: 2012-2021; grey) quagga mussel invasion (orange box: palatable; purple box: less-palatable species). Comparisons between the two decades were performed using zero-inflated GLMM (***: *p* < 0.001). Error bars represent one standard deviation from the mean (SD). For details see Supplementary Figure S5 and Tables S11 and S12.

The maximum summer water temperatures of Lake Müggelsee in the week before cyanobacteria sampling measured at the monitoring station in 0.5 m water depth, ranged between 16-30°C in the study years (Figure 4). The water temperature range of declining quagga mussel filtration rates, determined in our laboratory experiments (see Figure 1), was reached in several years before (2006 and 2010) and after the quagga mussel invasion (2013, 2014 and 2018). A comparison between littoral and pelagic water temperatures (measured at 1 m depth) showed that littoral water temperatures were up to 3°C (mean: 0.4°C) higher than pelagic water temperatures at the same depth, and the differences increased as temperature increased (Supplementary Figure S7).

**Figure 4:**
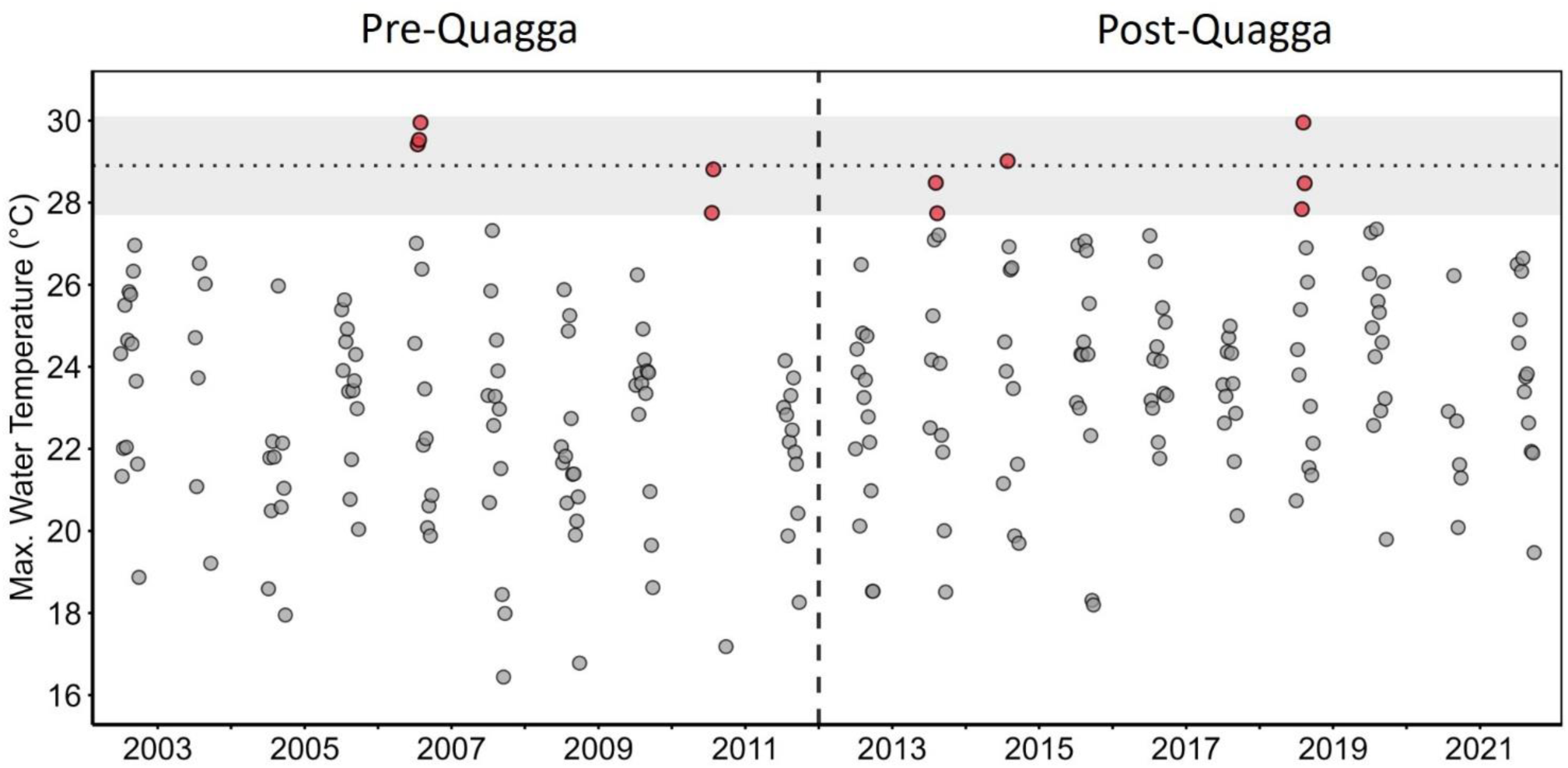
Summer (July–September) maximum water temperatures measured one week before phytoplankton samplings in Lake Müggelsee at a depth of 0.5 m, before (Pre-Quagga: 2002–2011; left) and after (Post-Quagga: 2012–2021; right) quagga mussel invasion. Red dots indicate values within the temperature range of declining quagga mussel filtration rates determined in the experiments shown in Figure 1 (dotted horizontal line at breakpoint 28.9°C; grey area represents 95% confidence interval: 27.7–30.1°C). The vertical dashed line marks the year of the quagga mussel invasion (2012). See Supplementary Figure S6 for details.

**Figure 5:**
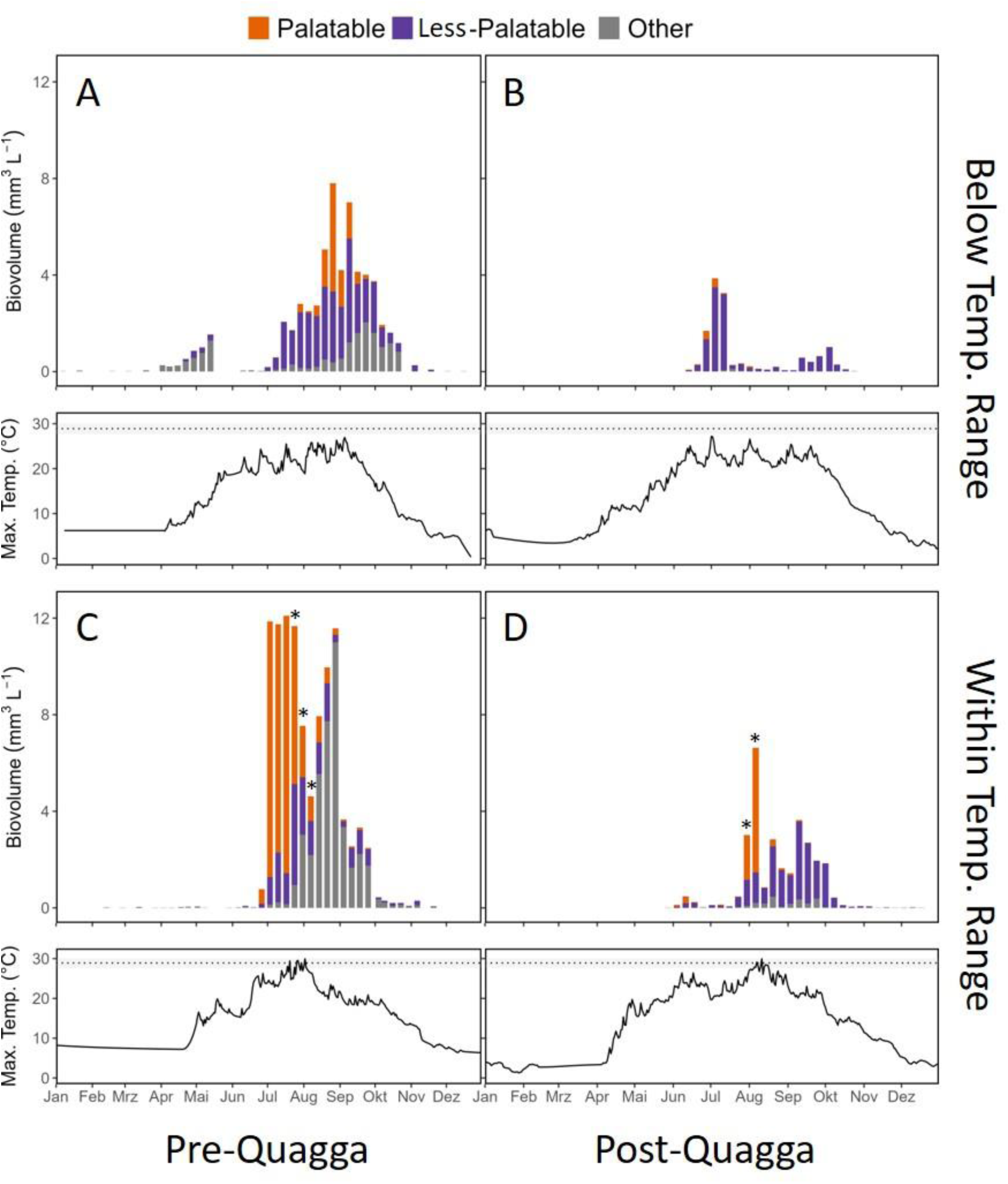
Biovolumes of palatable (orange, *Dolichospermum flos-aquae*), less-palatable (purple, *Anabaenopsis elenkinii*, *Aphanizomenon flos-aquae* and *Microcystis aeruginosa*) and non-classified cyanobacteria species (other, mainly *Planktothrix aghardii*) during selected years before. (Pre-Quagga; A: 2002, C: 2006) and after quagga mussel invasion (Post-Quagga; B: 2016, D: 2018). Examples were selected to represent years with daily maximum water temperatures reaching (C:2006, D: 2018) the temperature (temp.) range of declining quagga mussel filtration (see Figure 1) of 27.7-30.1 °C (grey box with BP at 28.9) and years with water temperatures below this range (A: 2002, B: 2016). Asterisks (*) indicate phytoplankton measurements at maximum daily temperatures within the critical water temperature range.

To serve as representative examples, the dynamics of the summer cyanobacteria bloom in Lake Müggelsee were analysed in more detail and compared for selected years with daily maximum water temperatures below or within the temperature range of declining quagga mussel filtration as determined in laboratory experiments (Figure 5A-D). Total cyanobacteria biovolumes were higher before (Figure 5A,C) than after quagga mussel invasion (Figure 5C,D; t-test: *p* < 0.001). In general, cyanobacteria blooms occurred when summer water temperatures were high. Palatable cyanobacteria formed blooms in both selected years before the quagga mussel invasion (Pre-Quagga, Figure 5A,C). However, this pattern differed in the years after quagga mussel invasion (Post-Quagga, Figure 5B,D). Here, high biovolumes of palatable cyanobacteria species only occurred when maximum daily lake temperatures reached the critical temperature range (Figure 5D). Palatable cyanobacteria almost completely disappeared after invasion, when summer water temperatures remained below this temperature range (Figure 5B).

Focusing on summer (July-September) data from 10 years before and after quagga mussel invasion, total cyanobacteria biovolume was significantly reduced after invasion (t-test: *p* < 0.001, Supplementary Figure S8). Considering the palatability of cyanobacteria, the biovolume of palatable cyanobacteria in Lake Müggelsee was significantly reduced following the quagga mussel invasion (GLMM: p < 0.001, Cohen’s d = –0.63, Figure 6 A,B). Only after the quagga mussel invasion segmented regression analysis and Davies test identified a significant BP of increasing palatable cyanobacteria biovolume above 27.7°C (95% CI: 26.9–28.4°C; p < 0.001). Less-palatable cyanobacteria showed no significant change in biovolume after invasion (GLMM: p = 0.345; Supplementary Tables S13 and S14; Figure S8), and no significant BP was detected.

**Figure 6:**
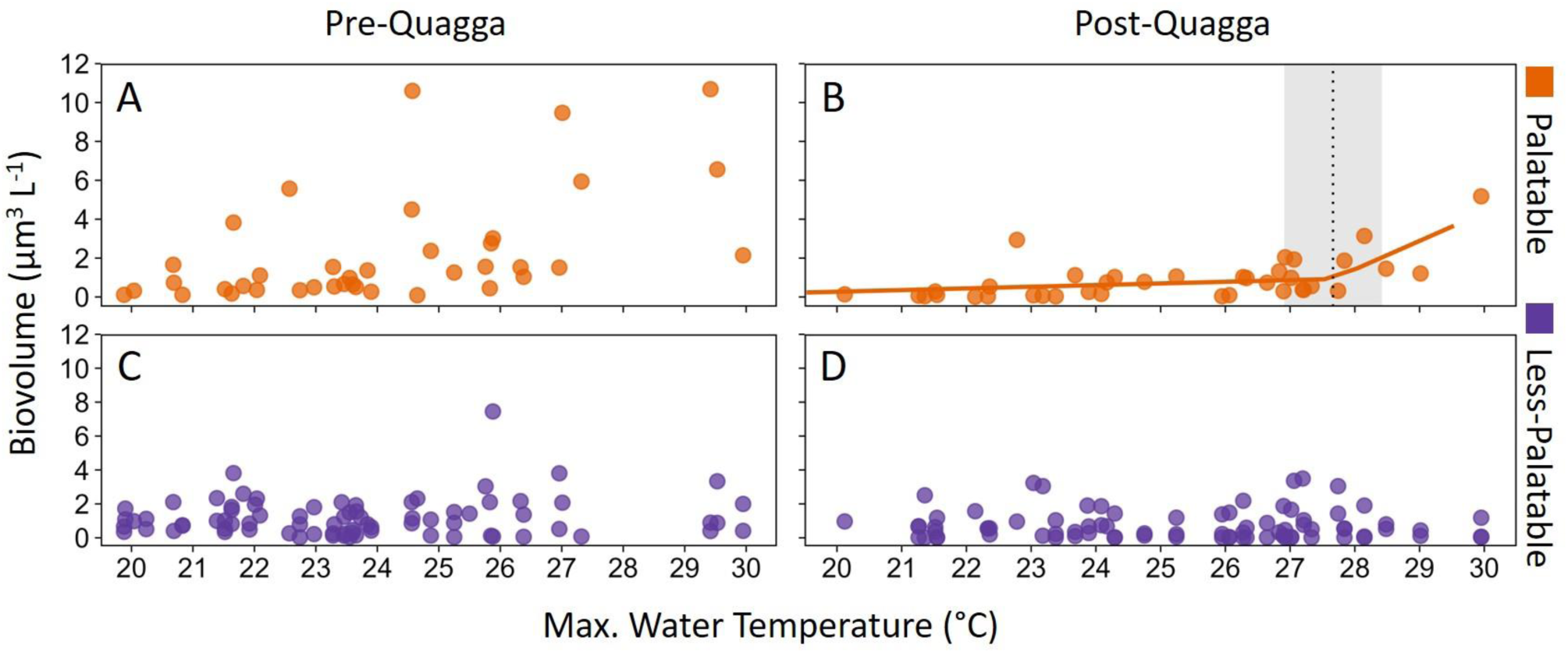
Summer (July-September) biovolumes of palatable (orange, top) and less-palatable (purple, bottom) cyanobacteria species in the decade before (Pre-Quagga: 2002-2011, left) and after. (Post-Quagga: 2012-2021, right) quagga mussel invasion in Lake Müggelsee along a gradient of maximum (Max.) water temperatures of hourly means at 0.5 m (measured in the week before the respective cyanobacteria bloom). The orange line shows segmented regression model including breakpoint (dotted black line) at 27.7°C (95% CI: 26.9–28.4°C) for palatable cyanobacteria after the quagga mussel invasion. No significant break point was observed in any of the other plots. For further details see Supplementary Tables S13-14 and Figures S9-10.

## 4. Discussion

Our findings support all four hypotheses. Laboratory experiments showed that quagga mussel filtration rates varied among typical bloom-forming cyanobacteria species. The potentially toxic, non-colony-forming species *Dolichospermum flos-aquae* was classified as palatable due to higher filtration rates of quagga mussels. In contrast, the potentially toxic, colony-forming species *Aphanizomenon flos-aquae, Anabaenopsis elenkinii* and *Microcystis aeruginosa* were classified as less-palatable cyanobacteria species for quagga mussels. No or very low filtration was measured above a temperature range of 27.7–30.1°C (95% CI with BP at 28.9°C) for three palatable species (*D. flos-aquae* and two green algae: *A. obliquus, C. vulgaris*), suggesting a critical upper thermal window for quagga mussel filtration. Analyses of long-term data from a temperate shallow lake 10 years before and after invasion support the hypothesis that quagga mussels can specifically reduce blooms of palatable toxic cyanobacteria such as *D. flos-aquae*. However, this effect is greatly reduced above a critical water temperature range of 26.9–28.4°C (95% CI with BP at 27.7°C), which confirms the results of the laboratory experiments. Therefore, quagga mussels can potentially control HABs when they are composed of palatable species and when water temperatures are below the critical temperature range for filtration. However, epilimnetic temperatures in lakes in the northern hemisphere invaded by quagga mussels will increasingly reach this critical temperature range in summer due to global warming. This will facilitate the occurrence of HABs in waters invaded by quagga mussels in two ways: by increasing the growth rates of cyanobacteria and by reducing the filtration effects of quagga mussels.

### 4.1. Differences in quagga mussel filtration on cyanobacteria species

Our results are consistent with those of previous studies on the relatively high filtration rates (palatability) of quagga mussels on *Dolichospermum flos-aquae* as compared to other cyanobacteria species (Knoll et al. 2008; Kirsch and Dzialowski 2012; Tang et al. 2014). Similarly, the filtration rates (palatability) on *Microcystis aeruginosa* were found to be rather low for both quagga mussels (Tang et al. 2014) and zebra mussels (Raikow et al. 2004; Vanderploeg et al. 2009). Information on dreissenid mussel filtration rates for the common cyanobacteria *Aphanizomenon flos-aquae* and *Anabaenopsis elenkinii* was unavailable. In our study, the filtration rates of quagga mussels on *Dolichospermum* and the green algae *Chlorella* and *Acutodesmus* at 24°C were similar to those reported in other studies (Dionisio Pires et al. 2004; Gopalakrishnan and Kashian 2020; Xia et al. 2021). Differences in filtration rates between the potentially toxic, non-colony-forming *D. flos-aquae* and the toxic, colony-forming *Microcystis aeruginosa* in our study support the notion from other studies that morphological traits such as colony formation are crucial in determining the palatability of phytoplankton for quagga mussels. However, other factors, such as unidentified cyanobacteria metabolites and other mechanisms may also be involved.

Our long-term lake data largely support the results of our laboratory experiments. We observed a reduction in palatable *Dolichospermum* biovolume after quagga mussel invasion, while the biovolume of all less-palatable cyanobacteria species remained statistically unchanged, although all means were numerically lower post-invasion. During the study period, P concentrations in the lake decreased, but P is not considered to be limiting cyanobacteria in Lake Müggelsee in summer (Shatwell and Köhler 2019). Instead, N limits cyanobacteria growth in Lake Müggelsee in summer, but DIN concentrations were not different for summers in the 10 years before and after the quagga mussel invasion. Although TN concentrations were different, particularly during summer, a large proportion of TN consists of refractory humic substances (see Figure 28 in Nixdorf et al. 2014), which are unavailable for phytoplankton growth (Fiedler et al. 2015) and therefore cannot explain the lower cyanobacteria biovolumes. The loss of phytoplankton-bound particulate N can be attributed to the filtration effects of the quagga mussels. The heterocyst frequency per biovolume of palatable *Dolichospermum* and less-palatable *Aphanizomenon* remained unchanged after quagga mussel invasion (Supplementary Figure S11 and Table S16). Additionally, no significant increase in heterocyst frequency in *Dolichospermum* occurred above the critical thermal window for mussel filtration (BP at 27.7°C identified in Figure 5), nor was there a significant linear relationship with temperature within these temperature segments (Supplementary Figure S12 and Tables S17-18). Therefore, we assume that the reduction in palatable cyanobacteria biovolume in Lake Müggelsee is due to the quagga mussel invasion rather than to changes in nutrient availability.

However, disentangling the effects of changes in nutrient loads and the quagga mussel invasion on lake phytoplankton dynamics still remains challenging. Including quagga mussel filtration in lake ecosystem models would be a useful tool, but the few studies using models that include dreissenid mussels have yet to disentangle these overlapping effects (Descy et al. 2003; Rowe et al. 2015; Woodruff et al. 2021).

### 4.2. Critical upper thermal window for quagga mussel filtration on cyanobacteria

Our laboratory experiments revealed a critical thermal window around 28°C, representing the boundary between active mussel filtration and thermal stress, in which filtration rates of quagga mussels sharply drop before they eventually die at 32°C. Thorp et al. (2002) suggested a sublethal temperature of 30°C for quagga mussels, and similar values have been reported for zebra mussels (Aldridge et al. 1995; Lei et al. 1996). Our analysis of long-term data from a temperate, shallow lake confirmed this critical thermal window. This has important implications for our understanding of the ecological impacts of quagga mussels. Within this thermal window, the control of cyanobacteria by mussels weakens dramatically, potentially allowing blooms to escape grazing control at higher temperatures. Additionally, quagga mussel die-offs during periods of extreme heat could release nutrients that may stimulate cyanobacterial growth. In this lake, quagga mussels were found at depths between 0-5.3 m (Wegner et al. 2019), and water temperatures in the deeper pelagic zones were about 1.3°C (mean summer Jul.-Sept. temperature difference to 5 m depth from 2002-2021, Supplementary Figure S10) lower than at a depth of 0.5 m. However, additional support for our critical thermal window comes from Quinn et al. (2014), who determined an upper temperature limit of 27.8°C based on the temperatures at 545 global occurrence points of quagga mussels, and used it for species distribution models.

Although prolonged exposure to high temperatures may lead to acclimation, others have found even lower lethal temperatures for quagga mussels (25°C), despite experimental acclimation (Spidle et al. 1995). Rising lake water temperatures are likely to be exacerbated in the near future, further stimulating the formation of HABs (Jöhnk et al. 2008; O’Neil et al. 2012; Visser et al. 2016), and the spread of invasive dreissenid mussels. Model predictions by Gallardo and Aldridge (2013) found strong benefits for zebra mussels under climate change scenarios and quagga mussels are expected to strongly benefit with thousands of susceptible lakes at risk of invasion (Gallardo and Aldridge 2025).

Our results show that quagga mussel filtration strongly reduces the biovolumes and therewith the bloom severity of the palatable cyanobacterium *Dolichospermum flos-aquae* below critical water temperatures. Blooms of this potentially toxic species are increasingly reported (Li et al. 2016; Sheik et al. 2022), and temperature-dependent reduction by quagga mussels can be relevant for their biocontrol. Less-palatable species such as *Microcystis aeruginosa* were filtered at lower rates (∼3 mL mg⁻¹ AFDW⁻¹ h⁻¹ at 24°C), but blooms may still be affected by quagga mussels (e.g. single cells or small colonies). The effectiveness of this process may depend on phenology, colony formation, and their total percentage within the phytoplankton community.

Cyanobacteria blooms reduced by quagga mussel filtration can return when water temperatures exceed the critical thermal window. Summer water temperatures in many lakes invaded by quagga mussels and those predicted to have suitable conditions (Quinn et al. 2014) already reach this thermal range, especially during heatwaves (Woolway et al. 2020; Dokulil et al. 2021). Quagga mussel grazing may no longer be able to control cyanobacteria growth, and in addition, certain cyanobacteria species grow particularly well in such warm conditions (Paerl and Huisman 2008). Increasing global warming is therefore expected to alter the interaction between two of the most significant stressors in northern hemisphere freshwaters, invasive quagga mussels and HABs, affecting freshwater ecosystem and human health (Griffith and Gobler 2020; Weyhenmeyer et al. 2024).

## Acknowledgments

This work was funded by the German Research Foundation (Deutsche Forschungsgemeinschaft, DFG) as part of the Research Training Group “Urban Water Interfaces” (GRK 2032/1). We gratefully acknowledge Katrin Preuss and Helga Täuscher (IGB) for conducting phytoplankton community analyses, and Julia Scheunert (IGB) for HPLC analyses. Field sampling was supported by Thomas Hintze, Jürgen Schreiber, Daniel Langenhaun, Tobias Goldhammer, and the IGB chemistry laboratory team as part of the institute’s long-term monitoring program for Lake Müggelsee. Experimental work benefited from the assistance of Zeno Mayr, Anna Schlegel, Anastasia Seelisch, Samantha Plank as well as mussel sampling by Lukas Dufner, and Renske Vroom. Renske as well co-supervised a student within this project. We thank Morgan Botrel and Misha Tseitlin for constructive discussions on statistical methodology and provision of code examples. Finally, we thank Thomas Mehner and participants of the IGB “Scientific Writing” workshop for stimulating discussions that improved this work. We gratefully acknowledge the two anonymous reviewers whose constructive comments strengthened the manuscript.

## Author contribution statement

JM and SH developed the hypotheses and designed the approach. JM, MEG and RN performed the laboratory experiments. JM, MEG and RN conducted analyses of the long-term data set. JK provided field data and information on cyanobacteria ecology in Lake Müggelsee. JM wrote the manuscript which was edited by all coauthors.

## Data Availability Statement

The data that support the findings of this study are available upon request from the corresponding author.

