## Supplementary material for "Blooms like it hot, but mussels do not: Influence of invasive quagga mussels on cyanobacteria during summer": Supllementary Material

**Orcid-ID: JM: https://orcid.org/0000-0003-1579-466X, JK: https://orcid.org/0000-0003-1894-2912, , SH: https://orcid.org/0000-0002-0585-2822**

*** Correspondence:**

Corresponding Author

### Content

#### Tables


- S1: Phytoplankton species, growth media, sampling data
- S2: Morphologies cyanobacteria
- S3: Microscopic countings (high)
- S4: Microscopic countings (low)
- S5: Filtration rate (descriptive statistics + CI)
- S6: Filtration rate GLM (species differences)
- S7: Filtration rate GLM Post hoc Tukey Pairwise (Effect size)
- S8: Filtration rate GLM for media
- S9: Nutrients - Bartlett test for homoscedasticity
- S10: Nutrients MANOVA
- S11: Cyanobacteria GLMM – Invasion on single species + interaction
- S12: Cyanobacteria GLMM – Invasion on single species + interaction (summary statistics)
- S13: Cyanobacteria GLMM – Invasion on palatable vs. less-palatable + interaction
- S14: Cyanobacteria GLMM – Invasion on palatable vs. less-palatable + interaction (summary statistics)
- S15: Cyanobacteria Breusch Pagan Homoscedasticity
- S16: Heterocyst formation of *Aphanizomenon* and *Dolichospermum* pre- vs. post invasion
- S17: Heterocyst formation of *Dolichospermum* above vs. below critical window
- S18: Heterocyst formation of *Dolichospermum* along maximum water temperature


#### Figures


- S1: Shell-length AFDW relationship
- S2: Filtration rate
- S3: Nutrients correlation
- S4: Nutrients details
- S5: Cyanobacteria details
- S6: Water temperature details
- S7: Pelagic-littoral temperature relationship
- S8: Cyanobacteria palatable vs. less-palatable biovolume
- S9 : Temperature biovolume homoscedasticity
- S10 : Comparison water temperatures at 0.5 and 5 m depth
- S11: Heterocyst formation of *Aphanizomenon* and *Dolichospermum*
- S12: Heterocyst formation of *Dolichospermum* along maximum water temperature

### 1 Supplementary Information Tables

Supplementary Table S1: Information on phytoplankton species, strain numbers, growth media used for culturing and date of quagga mussel sampling for the respective experiments. Strains were obtained from Culture Collection of Algae in Göttingen, Germany (SAG) and Pasteur Culture Collection of Cyanobacteria (PCC) in Paris, France or isolated from Lake Müggelsee. Information of toxins were derived from the algae collections. n.d. = not detected, - = not known to produce toxins. For general information on potential toxin production of the species, see Supplementary Table S2.

| Phytoplankton species | Strain number | Growth medium | Toxins | Quagga mussel sampling date |
|---|---|---|---|---|
| *Acutodesmus obliquus* (Turpin) (formerly *Scenedesmus obliquus)* | SAG 276-3a | Z4 | - | Jun. 22/29, Jul. 1, 2022 |
| *Chlorella vulgaris* Beijerinck | SAG 211.11j | Z8 | - | Dec. 6, 2022 |
| *Dolichospermum flos-aquae* (Lyngbye) | SAG 30.87 | Z8 | Anatoxin-a | Nov. 8, 2022 |
| *Anabaenopsis elenkinii* Miller | SAG 252.80 | BG11 + Spirulina medium (1:1) | n.d | Dec. 16, 2022 |
| *Aphanizomenon flos-aquae* (Linné) | Lake Müggelsee | Z8 | n.d | Sep. 15, 2023 |
| *Microcystis aeruginosa* Kützing | PCC 7820 | Z8 | Microcystins | Dec. 2, 2022 |
| Lake Müggelsee (high) | - | - | n.d | Aug. 23, 2023 |
| Lake Müggelsee (low) | - | - | n.d | Sep. 6, 2023 |

Supplementary Table S2: Morphologies and toxin formation in selected common bloom-forming cyanobacteria species used for filtration rate experiment with quagga mussels

| **Cyaobacterie species** | **Average size (cell/colony)** | **Shape** | **Colony-formation** | **Toxins** | **Cell structure** |
|---|---|---|---|---|---|
| ***Dolichospermum flos-aquae*** | Filament length (50-490 µm),<br><br>Filament width (~3.5 µm) (Huang et al. 2018; Gulati and Ejsmont-Karabin) | Filamentous structure (long chain of cells) (Xiao et al. 2016; Zhang et al. 2022) | Filamentous clumps (Tang et al. 2014; Li et al. 2016) | Microcystin, anatoxin-a (Van Der Merwe 2015; Li et al. 2016; Otero and Silva 2022) | Soft mucilage (some species) (Prasanna et al. 2006) |
| ***Microcystis aeruginosa*** | Single cells (~5 µm), Coloniues (up to 800 µm) (Aparicio Medrano et al. 2013; White and Sarnelle 2014) | Spheric cells (Dvořák et al. 2023) | Colony-forming (Tang et al. 2014; White and Sarnelle 2014) | Microcystin (Van Der Merwe 2015; Bui et al. 2018) | Colony surrounded by gelatinous sheath (Tang et al. 2014) |
| ***Anabaenopsis elenkinii*** | 3-6 µm (cell width) x 4-11 µm (cell length) (Komárek 2005; Aguilera et al. 2016) | Spheric to elongated multi-cell chains (Aguilera et al. 2016) | Filamentous, trichomes present (filaments solitary or in free clusters) (Aguilera et al. 2016) | Microcystin (Van Der Merwe 2015) | No sheath (Aguilera et al. 2016) |
| ***Aphanizomenon flos-aquae*** | Up to 2 cm (colony size) (Cirés and Ballot 2016; Lee et al. 2021) | Elongated multi-cell chains (Cirés and Ballot 2016; McGaraghan 2022) | Solitary trichomes or colony-forming filaments (Rekar and Hindák 2002; Cirés and Ballot 2016; McGaraghan 2022) | Anatoxin-a + saxitoxin (Cirés and Ballot 2016; McGaraghan 2022; Otero and Silva 2022) | No mucilage (Rekar and Hindák 2002) |

Supplementary Table S3: Microscopic countings and conversion into biovolume and share of total sample biovolume for Lake Müggelsee (high) samples.

| **Taxon** | | | **Individuals** | **Biovolume** | **Share of tot. Biov.** |
|---|---|---|---|---|---|
| **Genus** | **Species** | **Class** | **(Ind. $L^{-1}$)** | **($\mu m^3$ $L^{-1}$)** | **(%)** |
| *Microcystis* | *spp.* | Cyanobacteria | 890118 | 1.4 | 10.8 |
| *Aphanizomenon* | *flos aquae* | Cyanobacteria | 366519 | 0.5 | 3.6 |
| *Microcystis* | *aeruginosa* | Cyanobacteria | 314159 | 7.3 | 57.2 |
| *Pseudanabaena* | *mucicola* | Cyanobacteria | 104720 | 0.1 | 0.5 |
| *Pseudanabaena* | *mucicola* | Cyanobacteria | 785398 | 0.0 | 0.1 |
| *Cyanobacteria* | Round single cells | Cyanobacteria | 26703538 | 1.0 | 8.1 |
| *Ankyra* | *judayii* | Chlorophyta | 589049 | 0.0 | 0.0 |
| *Closterium* | *aciculare* | Charophyta | 8000 | 0.1 | 0.6 |
| *Chrysoflagellata* | *spp.* | Chrysophyta | 1767146 | 0.1 | 0.4 |
| *Cryptomonas* | *spp.* | Cryptophyceae | 680678 | 0.3 | 2.4 |
| *Cryptomonas* | *spp.* | Cryptophyceae | 314159 | 0.6 | 5.0 |
| *Cryptomonas* | *spp.* | Cryptophyceae | 916298 | 1.0 | 7.5 |
| *Rhodomonas* | *minuta/lacustris* | Cryptophyceae | 4280420 | 0.1 | 0.7 |
| *Ceratium* | *hirundinella* | Dinophyceae | 4000 | 0.2 | 1.8 |
| *Nitzschia* | *fonticola* | Diatoms | 52360 | 0.0 | 0.3 |
| *Diatoms* | *spp.* | Diatoms | 589049 | 0.0 | 0.4 |
| *Nitzschia* | *fonticola* | Diatoms | 785398 | 0.1 | 0.6 |

Supplementary Table S4: Microscopic countings and conversion into biovolume and share of total sample biovolume (BV) for Lake Müggelsee (low) samples.

| **Taxon** | | | **Individuals** | **Biovolume** | **Share of total BV** |
|---|---|---|---|---|---|
| **Genus** | **Species** | **Class** | **(Ind. $L^{-1}$)** | **($\mu m^3$ $L^{-1}$)** | **(%)** |
| *Dolichospermum* | *affine* | Cyanobacteria | 471239 | 0.3 | 1.4 |
| *Dolichospermum* | *circinale* | Cyanobacteria | 78540 | 0.1 | 0.4 |
| *Dolichospermum* | *flos-aquae* | Cyanobacteria | 471239 | 0.3 | 1.4 |
| *Microcystis* | *spp.* | Cyanobacteria | 549779 | 1.1 | 5.1 |
| *Microcystis* | *aeruginosa* | Cyanobacteria | 903208 | 7.5 | 33.8 |
| *Pseudanabaena* | *Mucicola* | Cyanobacteria | 157080 | 0.1 | 0.3 |
| *Aphanizomenon* | *flos-aquae* | Cyanobacteria | 12252211 | 9.7 | 43.5 |
| *Aphanizomenon* | *issatschenkoi* | Cyanobacteria | 157080 | 0.1 | 0.6 |
| *Pseudanabaena* | *spp.* | Cyanobacteria | 39270 | 0.0 | 0.0 |
| *Dolichospermum* | *mendotae* | Cyanobacteria | 78540 | 0.0 | 0.0 |
| *Pseudanabaena* | *mucicola* | Cyanobacteria | 1374447 | 0.0 | 0.1 |
| *Cyanobacteria* | Round single cells | Cyanobacteria | 44767696 | 1.7 | 7.8 |
| *Chrysoflagellata* | *spp.* | Chrysophyta | 589049 | 0.0 | 0.2 |
| *Chrysochromulina* | *spp.* | Chrysophyta | 589049 | 0.0 | 0.1 |
| *Cryptomonas* | *spp.* | Cryptophyceae | 78540 | 0.0 | 0.2 |
| *Ceratium* | *furcoides* | Dinophyceae | 8000 | 0.5 | 2.1 |
| *Ceratium* | *hirundinella* | Dinophyceae | 4000 | 0.2 | 0.9 |
| *Nitzschia* | *fonticola* | Diatoms | 235619 | 0.1 | 0.4 |
| *Fragilaria* | *angustissima* | Diatoms | 39270 | 0.2 | 1.0 |
| *Nitzschia* | *fonticola* | Diatoms | 1178097 | 0.1 | 0.6 |

Supplementary Table S5: Cyanobacterial filtration rates by quagga mussels across experimental temperatures. Mean filtration rates, standard deviations (SD), standard errors (SE), 95% confidence intervals (CI), and 95% confidence intervals ($CI_{lower}$, $CI_{upper}$) for four cyanobacterial species (*Dolichospermum*, *Anabaenopsis*, *Microcystis*, and *Aphanizomenon*) at five experimental temperatures (24, 26, 28, 30, 32°C)

| Genus | Temp | n | mean | sd | se | $ci_{lower}$ | $ci_{upper}$ |
|---|---|---|---|---|---|---|---|
| | 24 | 4 | 3.02 | 0.59 | 0.30 | 2.44 | 3.60 |
| | 26 | 4 | 4.00 | 1.62 | 0.81 | 2.41 | 5.58 |
| ***Dolichospermum*** | 28 | 4 | 4.26 | 1.30 | 0.65 | 2.99 | 5.53 |
| | 30 | 4 | 3.23 | 0.58 | 0.29 | 2.67 | 3.80 |
| | 32 | 4 | 0.60 | 0.56 | 0.28 | 0.05 | 1.15 |
| | 24 | 4 | -0.16 | 0.98 | 0.49 | -1.12 | 0.80 |
| | 26 | 4 | 0.39 | 1.33 | 0.67 | -0.91 | 1.70 |
| ***Aphanizomenon*** | 28 | 4 | -0.29 | 1.08 | 0.54 | -1.35 | 0.77 |
| | 30 | 4 | 0.32 | 0.38 | 0.19 | -0.05 | 0.70 |
| | 32 | 4 | 0.00 | 0.95 | 0.47 | -0.93 | 0.93 |
| | 24 | 4 | 2.31 | 0.95 | 0.47 | 1.38 | 3.24 |
| | 26 | 4 | -0.20 | 0.34 | 0.17 | -0.53 | 0.14 |
| ***Anabaenopsis*** | 28 | 4 | 0.38 | 0.50 | 0.25 | -0.11 | 0.88 |
| | 30 | 4 | 1.44 | 0.70 | 0.35 | 0.76 | 2.12 |
| | 32 | 4 | 0.59 | 0.44 | 0.22 | 0.15 | 1.02 |
| | 24 | 4 | 2.68 | 0.95 | 0.48 | 1.75 | 3.62 |
| | 26 | 4 | 1.48 | 0.89 | 0.44 | 0.61 | 2.35 |
| ***Microcystis*** | 28 | 4 | 1.03 | 1.72 | 0.86 | -0.66 | 2.71 |
| | 30 | 4 | -0.27 | 0.60 | 0.30 | -0.86 | 0.32 |
| | 32 | 4 | -0.25 | 0.99 | 0.50 | -1.22 | 0.72 |

Supplementary Table S6: Summary of General Linear Model (GLM) results examining the effects of cyanobacterial genus and water temperature on quagga mussel filtration rates. Fixed effects include genus (*Dolichospermum*, *Anabaenopsis*, *Microcystis*, *Aphanizomenon*), temperature (24, 26, 28, 30, 32°C), and their interaction. Columns show degrees of freedom (df), Type III sum of squares test statistic, p-values ($\alpha = 0.05$), and effect sizes ($\eta^2$ = eta-squared, partial eta-squared).

| **Effect** | **df** | **$\chi^2$** | **p-value** | **$\eta^2$** |
|---|---|---|---|---|
| Genus | 3 | 27.64 | <0.001 | 0.11 |
| Temperature | 4 | 37.03 | <0.001 | 0.14 |
| Genus × Temperature | 12 | 55.09 | <0.001 | 0.20 |

Supplementary Table S7: Post-hoc Tukey pairwise contrasts with effect sizes comparing filtration rate differences between all cyanobacterial species (*Dolichospermum* vs. *Aphanizomenon*, *Anabaenopsis*, and *Microcystis*) across five experimental water temperatures (24, 26, 28, 30, 32°C). Columns show the estimated difference in filtration rate between species pairs (estimate), standard error (SE), t-ratio test statistic, degrees of freedom (df), p-value (adjusted for multiple comparisons), and Cohen's d effect size.

| contrast | Temp | estimate | SE | df | t.ratio | p.value | cohen d |
|---|---|---|---|---|---|---|---|
| *Anabaenopsis - Microcystis* | 24 | 3.18 | 0.67 | 60 | 4.72 | 0.00 | 3.83 |
| | 26 | 0.71 | 0.67 | 60 | 1.05 | 0.72 | 0.85 |
| | 28 | 0.34 | 0.67 | 60 | 0.50 | 0.96 | 0.41 |
| | 30 | -2.48 | 0.67 | 60 | -3.67 | 0.00 | -2.98 |
| | 32 | -2.85 | 0.67 | 60 | -4.22 | 0.00 | -3.43 |
| *Aphanizomenon - Anabaenopsis* | 24 | -0.37 | 0.67 | 60 | -0.55 | 0.95 | -0.44 |
| | 26 | 3.60 | 0.67 | 60 | 5.35 | 0.00 | 4.34 |
| | 28 | 4.19 | 0.67 | 60 | 6.22 | 0.00 | 5.05 |
| | 30 | 2.52 | 0.67 | 60 | 3.73 | 0.00 | 3.03 |
| | 32 | 0.59 | 0.67 | 60 | 0.87 | 0.82 | 0.71 |
| *Aphanizomenon - Microcystis* | 24 | -1.09 | 0.67 | 60 | -1.61 | 0.38 | -1.31 |
| | 26 | -1.68 | 0.67 | 60 | -2.49 | 0.07 | -2.02 |
| | 28 | 4.55 | 0.67 | 60 | 6.75 | 0.00 | 5.48 |
| | 30 | 3.88 | 0.67 | 60 | 5.76 | 0.00 | 4.67 |
| | 32 | 3.24 | 0.67 | 60 | 4.80 | 0.00 | 3.90 |
| *Dolichospermum - Anabaenopsis* | 24 | -0.67 | 0.67 | 60 | -1.00 | 0.75 | -0.81 |
| | 26 | -1.31 | 0.67 | 60 | -1.95 | 0.22 | -1.58 |
| | 28 | -0.64 | 0.67 | 60 | -0.95 | 0.78 | -0.77 |
| | 30 | 2.91 | 0.67 | 60 | 4.32 | 0.00 | 3.50 |
| | 32 | 1.79 | 0.67 | 60 | 2.66 | 0.05 | 2.16 |
| *Dolichospermum - Aphanizomenon* | 24 | 3.51 | 0.67 | 60 | 5.20 | 0.00 | 4.22 |
| | 26 | -1.12 | 0.67 | 60 | -1.66 | 0.36 | -1.34 |
| | 28 | 0.60 | 0.67 | 60 | 0.89 | 0.81 | 0.72 |
| | 30 | 1.71 | 0.67 | 60 | 2.54 | 0.06 | 2.06 |
| | 32 | 0.60 | 0.67 | 60 | 0.89 | 0.81 | 0.72 |
| *Dolichospermum - Microcystis* | 24 | 0.01 | 0.67 | 60 | 0.02 | 1.00 | 0.02 |
| | 26 | 0.85 | 0.67 | 60 | 1.26 | 0.59 | 1.02 |
| | 28 | -0.59 | 0.67 | 60 | -0.87 | 0.82 | -0.70 |

| | | | | | | |
|---|---|---|---|---|---|---|
| 30 | 0.25 | 0.67 | 60 | 0.37 | 0.98 | 0.30 |
| 32 | 0.84 | 0.67 | 60 | 1.24 | 0.60 | 1.01 |

Supplementary Table S8: Type III ANOVA results of generalized linear model testing culture medium type (Supplementary Table S1) effects on cyanobacterial filtration rates across temperatures 24–32°C.

| **Effect** | **Sum of Squares** | **df** | **F-value** | **p-value** |
|---|---|---|---|---|
| Media Type | 0.654 | 1 | 0.255 | 0.615 |
| Temperature | 15.421 | 4 | 1.501 | 0.211 |
| Media Type × Temperature | 17.723 | 4 | 1.725 | 0.154 |
| Residuals | 179.852 | 70 | — | — |

Supplementary Table S9: Summary of Bartlett test statistics for log-transformed nutrient values separated by invasion status as test for homoscedasticity. DF: Degrees of freedom

| Nutrient | Bartlett's K-squared | DF | p-value | Result |
|---|---|---|---|---|
| srp | 0.62 | 1 | 0.432 | homoscedasticity |
| tp | 0.06 | 1 | 0.811 | homoscedasticity |
| din | 2.33 | 1 | 0.127 | homoscedasticity |
| tn | 2.13 | 1 | 0.144 | homoscedasticity |

Supplementary Table S10: Multivariate Analysis of Variance (MANOVA) results testing whether nutrient concentrations for dissolved inorganic nitrogen (DIN), soluble reactive phosphorus (SRP), total nitrogen (TN), and total phosphorus (TP) differed between pre-invasion (2002-2011) and post-invasion (2012-2021) periods in Lake Müggelsee.

| Nutrient | F_statistic | p_value | eta_squared | Pre_mean | Post_mean | Percent_change |
|---|---|---|---|---|---|---|
| DIN | 0.27 | 0.60 | 0.001 | 0.07 | 0.08 | 9.38 |
| SRP | 63.25 | < 0.001 | 0.21 | 100.72 | 41.31 | -58.98 |
| TN | 61.52 | < 0.001 | 0.21 | 1.015 | 0.81 | -19.94 |
| TP | 156.23 | < 0.001 | 0.40 | 232.67 | 106.36 | -54.29 |

Supplementary Table S11: Fixed effects coefficients from the conditional component of the zero-inflated Gamma GLMM examining the influence of quagga mussel invasion and cyanobacteria genus on cyanobacteria biovolume (BV). The model included the invasion status Pre-Quagga (2002-2011) and Post-Quagga (2012-2021), genus (*Dolichospermum*, *Anabaenopsis*, *Aphanizomenon*, and *Microcystis*), and their interaction as fixed effects, with a random intercept for sampling date. *Dolichospermum* under Post-Quagga conditions serves as the reference level. Estimate: Log-scale coefficient estimate from the GLMM; Std. Error: Standard error of the coefficient estimate; z-value: Test statistic (estimate divided by standard error); Exp (Estimate): Exponentiated coefficient, representing the multiplicative effect on the response variable (BV). Significance levels: *** p<0.001.

| Term | Estimate | Std. Error | z-value | p-value | Exp (Estimate) | Interpretation |
|---|---|---|---|---|---|---|
| (Intercept) | -1.45022 | 0.17622 | -8.230 | < 0.001 *** | 0.235 | Baseline log-biovolume (for *Dolichospermum* under Post-Quagga conditions) |
| Pre-Quagga | 1.23502 | 0.25710 | 4.804 | < 0.001 *** | 3.438 | Main effect of Pre-Quagga on *Dolichospermum*: biovolume increases by about 3.44× relative to Post-Quagga |
| *Anabaenopsis* | -2.12474 | 0.29068 | -7.310 | < 0.001 *** | 0.119 | Baseline difference: *Anabaenopsis* has lower biovolume than *Dolichospermum* under Post-Quagga conditions |
| *Aphanizomenon* | 0.18048 | 0.17897 | 1.008 | 0.313 | 1.198 | Baseline difference: *Aphanizomenon* biovolume is ~1.2× that of *Dolichospermum* under Post-Quagga (not significant) |
| *Microcystis* | -0.04526 | 0.20936 | -0.216 | 0.829 | 0.956 | Baseline difference: *Microcystis* biovolume is nearly identical to that of *Dolichospermum* under Post-Quagga |
| Pre-Quagga × *Anabaenopsis* | 0.64478 | 0.47001 | 1.372 | 0.170 | 1.906 | Additional invasion effect for *Anabaenopsis* (not statistically significant) |
| Pre-Quagga × *Aphanizomenon* | -0.32036 | 0.27098 | -1.182 | 0.237 | 0.726 | Additional invasion effect for *Aphanizomenon* (not statistically significant) |
| Pre-Quagga × *Microcystis* | -0.26520 | 0.36476 | -0.727 | 0.467 | 0.767 | Additional invasion effect for *Microcystis* (not statistically significant) |

Supplementary Table S12: Summary statistics for the zero-inflated Gamma GLMM assessing the effects of Quagga mussel invasion and genus on cyanobacteria biovolume. The model was fitted with a log-link function and included a random intercept for sampling date to account for temporal autocorrelation. AIC: Akaike Information Criterion; BIC: Bayesian Information Criterion.

| **Statistic** | **Value** |
|---|---|
| AIC | 174.1 |
| BIC | 251.7 |
| Log-Likelihood | -69.1 |
| Conditional Deviance | 138.1 |
| Residual Degrees of Freedom | 531 |
| Random Effect Variance (Date) | 0.924 |
| Dispersion Parameter ($\sigma^2$) | 1.24 |
| Number of Observations | 549 |
| Number of Dates (Groups) | 257 |

Supplementary Table S13: Summary statistics for the zero-inflated Gamma GLMM assessing the effects of quagga mussel invasion and palatable and less-palatable groups on cyanobacteria biovolume. The model was fitted with a log-link function and included a random intercept for sampling date to account for temporal autocorrelation. AIC: Akaike Information Criterion; BIC: Bayesian Information Criterion.

| **Parameter** | **Estimate** | **Std. Error** | **z value** | **p-value** | **Exp (Estimate)** | **Interpretation** |
|---|---|---|---|---|---|---|
| Intercept | -1.333 | 0.177 | -7.549 | <0.001*** | 0.264 | Baseline log-biovolume (Palatable, Post-Quagga) |
| Pre-Quagga | 1.273 | 0.260 | 4.900 | <0.001*** | 3.572 | Palatable taxa biovolume was 3.57× higher pre-invasion |
| Less-Palatable | -0.099 | 0.172 | -0.577 | 0.564 | 0.905 | No significant difference between palatability groups post-invasion |
| Pre-Quagga × Less-Palatable | -0.253 | 0.268 | -0.944 | 0.345 | 0.776 | Non-significant difference in invasion effect between palatability groups |

Supplementary Table S14: Summary statistics for the zero-inflated Gamma GLMM assessing the effects of quagga mussel invasion and palatable and less-palatable groups on cyanobacteria biovolume. The model was fitted with a log-link function and included a random intercept for sampling date to account for temporal autocorrelation. AIC: Akaike Information Criterion; BIC: Bayesian Information Criterion.

| **Statistic** | **Value** |
|---|---|
| AIC | 214.1 |
| BIC | 257.2 |
| Log-Likelihood | -97.1 |
| Random Effect Variance (Date) | 0.715 |
| Dispersion Parameter ($\sigma^2$) | 1.430 |
| Number of Observations | 549 |
| Number of Dates (Groups) | 257 |

Supplementary Table S15: Results of the Breusch-Pagan test examining the homoscedasticity assumption in the linear regression model of cyanobacterial biovolume as a function of maximum-water temperature. The test statistic (BP) follows a chi-square distribution with the specified degrees of freedom (DF). The p-value indicates the probability of observing the given test statistic or more extreme values under the null hypothesis of homoscedasticity. A p-value greater than 0.05 suggests that the assumption of constant error variance is satisfied.

| Cyano-Type | Invasion | BP | DF | p-value | Result |
|---|---|---|---|---|---|
| Palatable | Pre-Quagga | 3.62 | 1 | 0.06 | homoscedasticity |
| Palatable | Post-Quagga | 2.9403 | 1 | 0.09 | homoscedasticity |
| Less-Palatable | Pre-Quagga | 1.2462 | 1 | 0.26 | homoscedasticity |
| Less-Palatable | Post-Quagga | 0.29084 | 1 | 0.59 | homoscedasticity |

Supplementary Table S16: Results of two-sample t-tests comparing pre- and post-quagga periods for biovolume, heterocyst density per litre, and heterocyst density per biovolume (BV) of *Aphanizomenon* and *Dolichospermum*. Reported are t-statistics and associated p-values for each genus–variable combination.

| Genus | Variable | t_stat | p_value |
|---|---|---|---|
| *Aphanizomenon* | Biovolume (BV) | 4.14 | < 0.001 |
| *Aphanizomenon* | Heterocysts per L | 3.30 | 0.001 |
| *Aphanizomenon* | Heterocysts per BV | 2.42 | 0.016 |
| *Dolichospermum* | Biovolume (BV) | 3.12 | 0.002 |
| *Dolichospermum* | Heterocysts per L | 2.93 | 0.004 |
| *Dolichospermum* | Heterocysts per BV | 0.50 | 0.614 |

Supplementary Table S17: Wilcoxon rank-sum tests comparing mean heterocyst frequencies per biovolume for *Dolichospermum* above (≥27.7°C) vs. below (<27.7°C) the critical thermal threshold from long-term data analysis in Figure 6, separated by pre- and post-quagga periods. Reported are sample sizes, means, and p-values.

| Period | n_total | n_below | n_above | mean_below | mean_above | wilcox_p | signif |
|---|---|---|---|---|---|---|---|
| Pre-Quagga | 96 | 91 | 5 | 280882.13 | 195006.16 | 0.89 | n.s. |
| Post-Quagga | 109 | 103 | 6 | 242059.87 | 298656.70 | 0.26 | n.s. |

Supplementary Table S18: Linear regression slopes of heterocyst frequency per biovolume of *Dolichospermum* along maximum (Max.) temperature, segmented above (≥27.7°C) vs. below (<27.7°C) the critical thermal threshold identified in Figure 6, for pre- and post-quagga periods separately. Reported are slopes, p-values, and sample sizes per segment.

| Period | segment | slope | p | n |
|---|---|---|---|---|
| Pre-Quagga | below | 559.1 | 0.979 | 91 |
| Pre-Quagga | above | 162412.5 | 0.168 | 5 |
| Post-Quagga | below | 31527.7 | 0.098 | 103 |
| Post-Quagga | above | -123605.5 | 0.198 | 6 |

### 2 Supplementary Figures

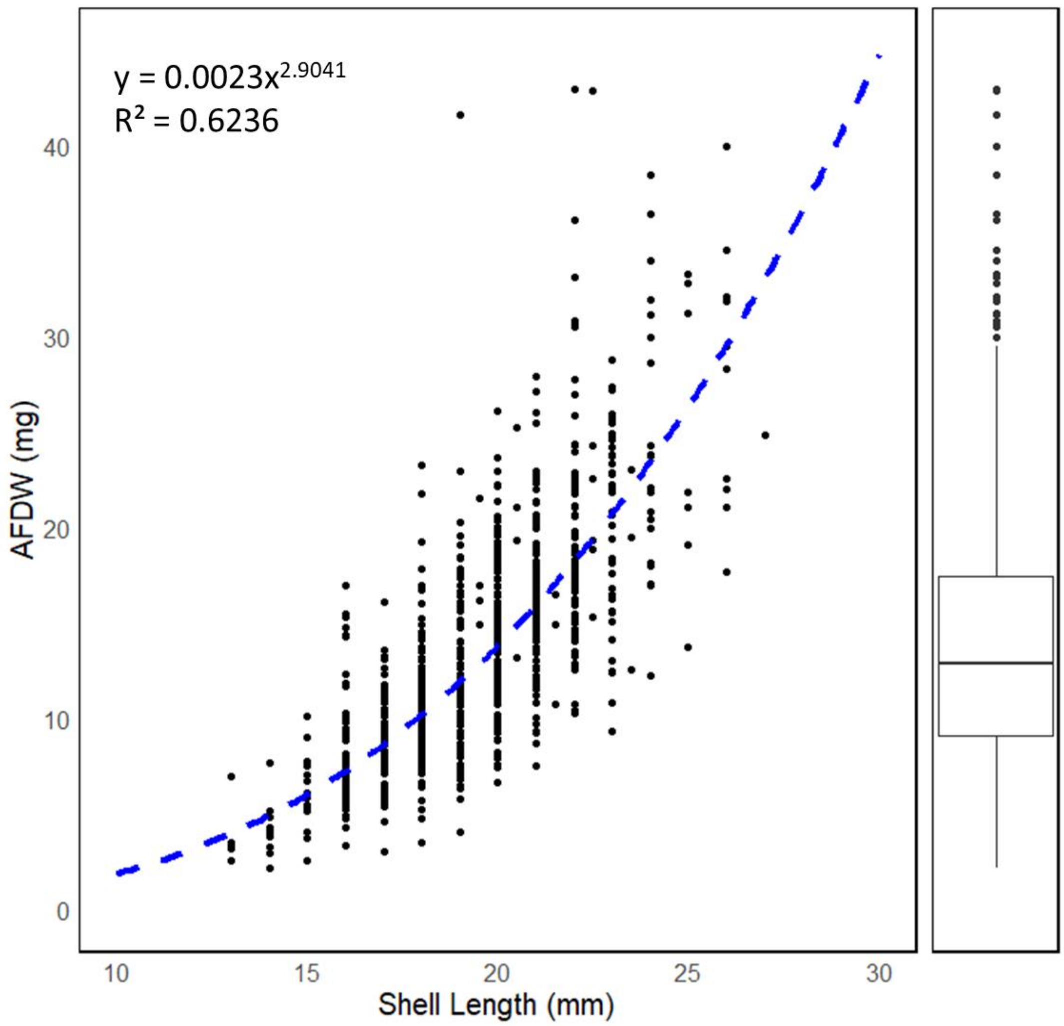


Supplementary Figure S1: Quagga mussel shell length to ash-free dry weight (AFDW) relationship (n = 827) from filtration rate experiments including regression.

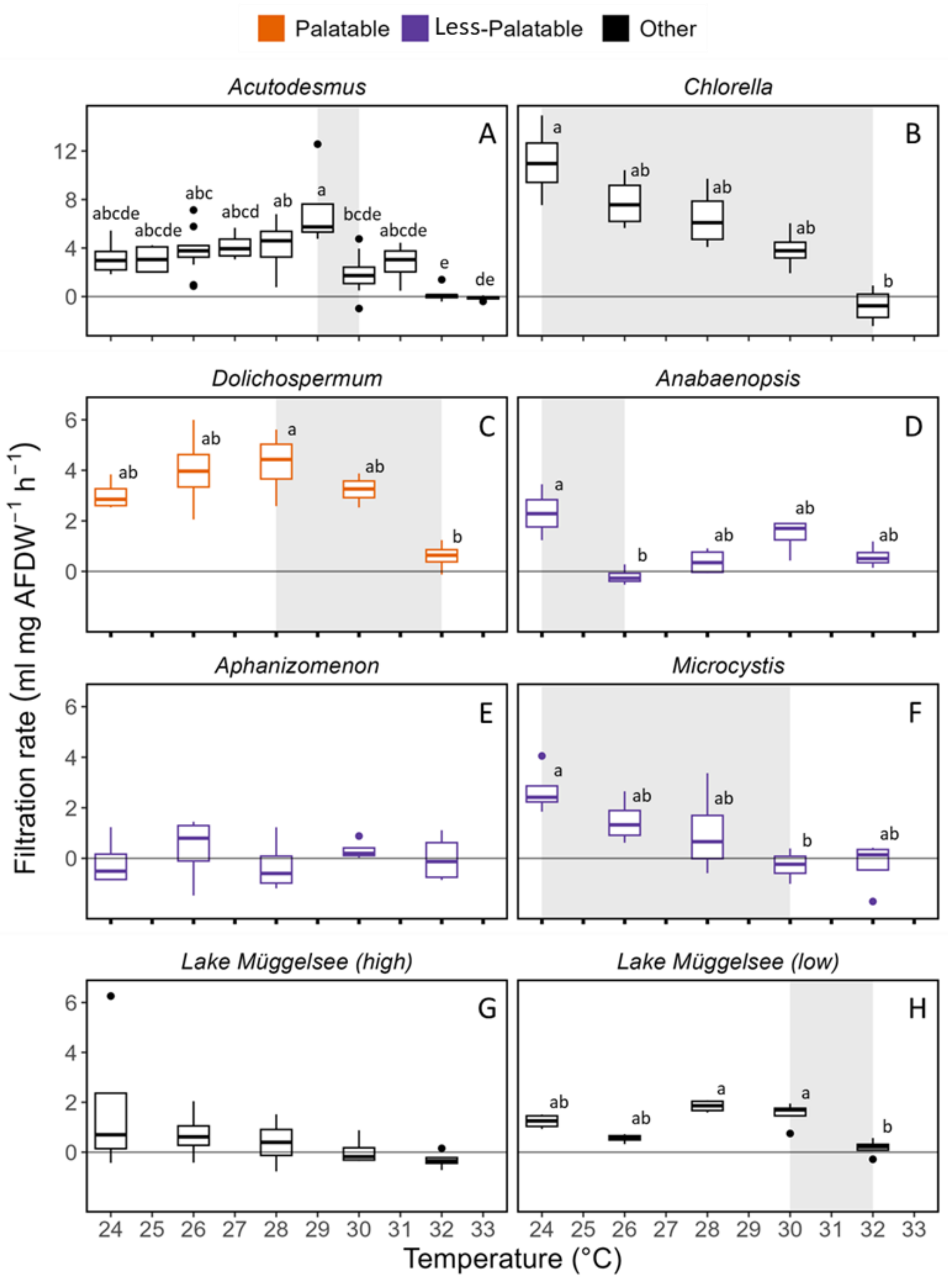


Supplementary Figure S2: Filtration rates of quagga mussels on different phytoplankton species (green algae: *Acutodesmus obliquus* (A) and *Chlorella vulgaris* (B); cyanobacteria: *Dolichospermum flos-aquae* (C), *Anabaenopsis elenkinii* (D), *Aphanizomenon flos-aquae* (E) and *Microcystis aeruginosa* (F)) and two mixed phytoplankton samples from Lake Müggelsee containing high concentrations (G)

(chlorophyll *a*: ~30 µg $L^{-1}$, 95% cyanobacteria, mainly *M. aeruginosa*) and low concentrations (H) (chlorophyll *a*: ~ 15 µg $L^{-1}$, 80% cyanobacteria, mainly *M.* aeruginosa) at different water temperatures (24–32°C). Boxplots show median, quartiles, and outliers.

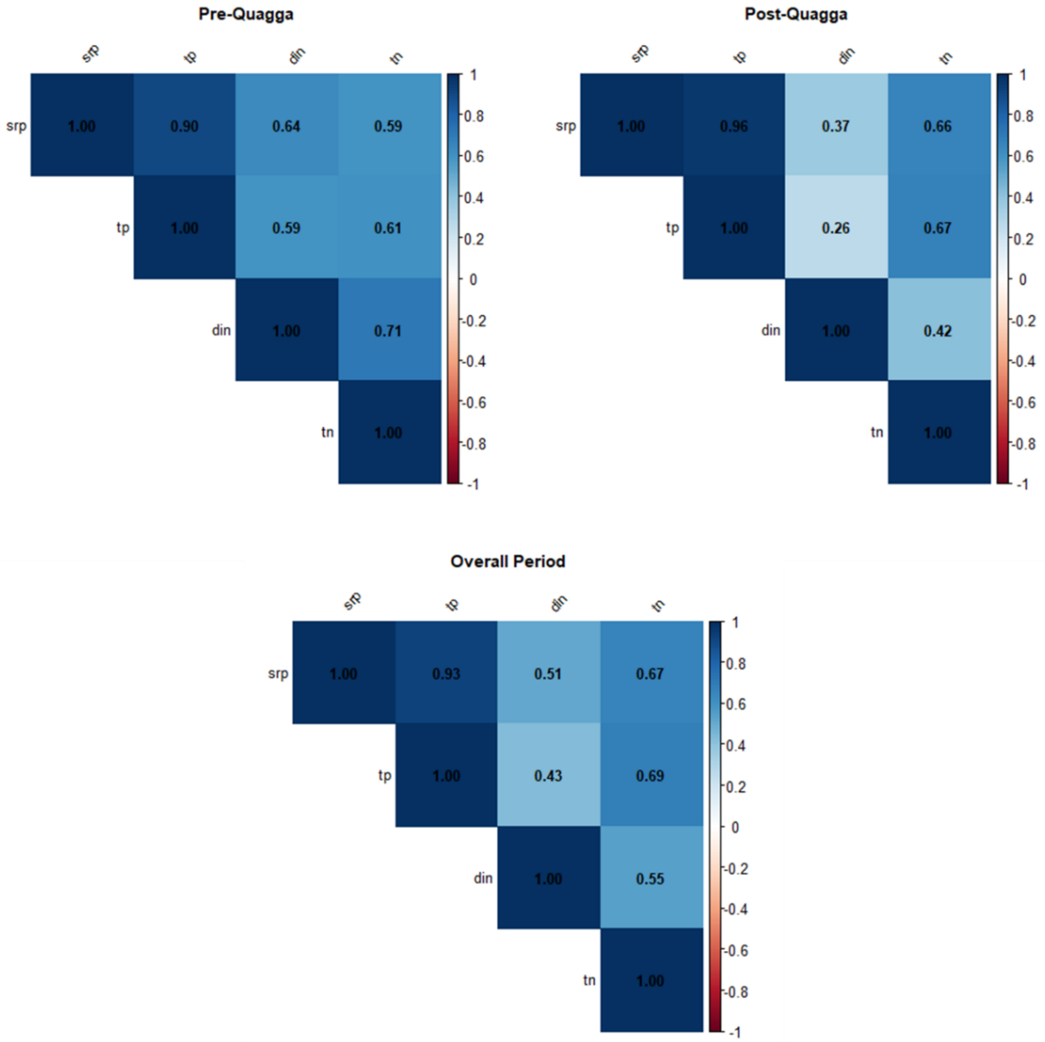


Supplementary

Figure S3: Pearson-correlation matrices of summer (July-September) nutrient concentrations (DIN: dissolved inorganic nitrogen, SRP: soluble reactive phosphorus, TN: total nitrogen, TP: total phosphorus) in Lake Müggelsee before (2002-2011), after (2012-2021) quagga mussel invasion and for the whole period (2002-2021). Color intensity and size of the correlation coefficients indicate the strength of correlation, with blue representing positive correlations and red representing negative correlations. All correlations were calculated using complete observations only.

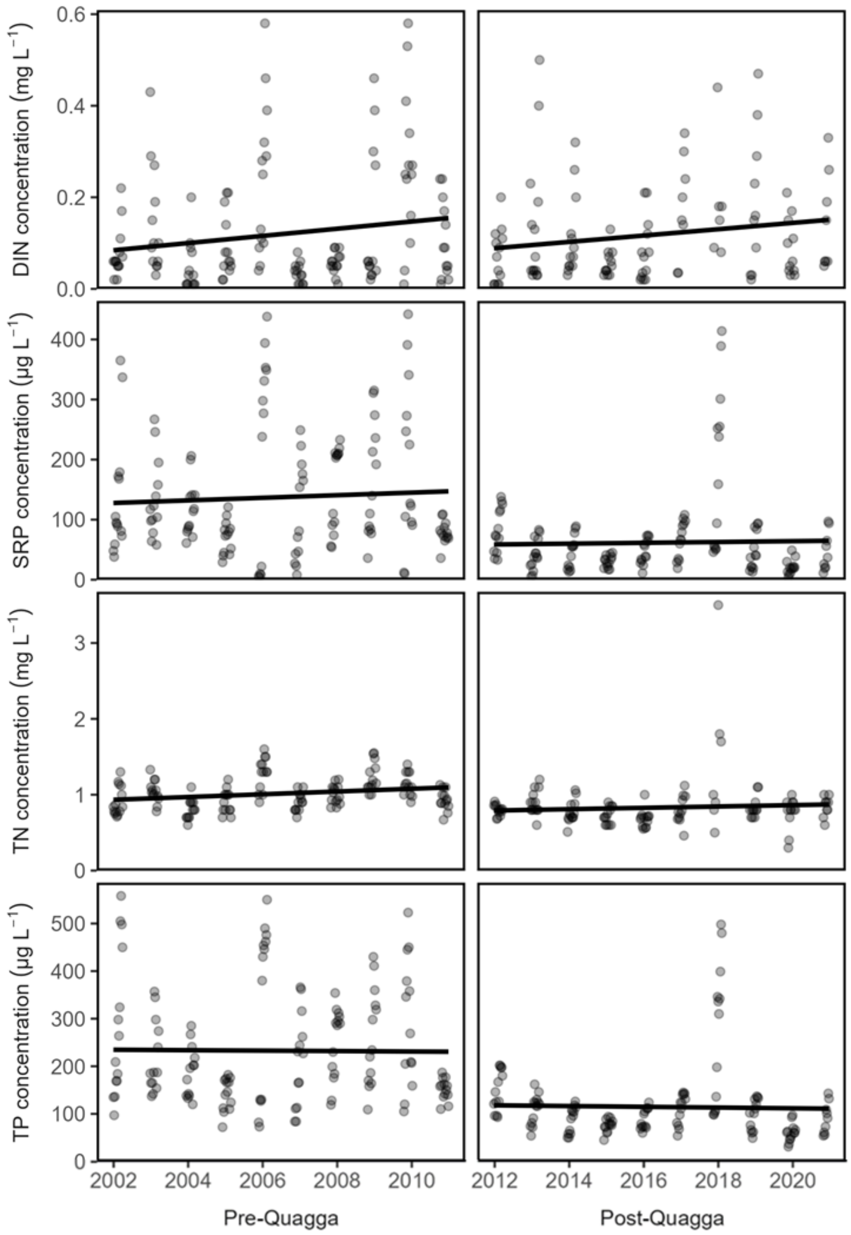

Supplementary Figure S4: Summer (July-September) nutrient concentrations (DIN: dissolved inorganic nitrogen, SRP: soluble reactive phosphorus, TN: total nitrogen, TP: total phosphorus) in Lake Müggelsee before (2002-2011) and after (2012-2021) quagga mussel invasion.

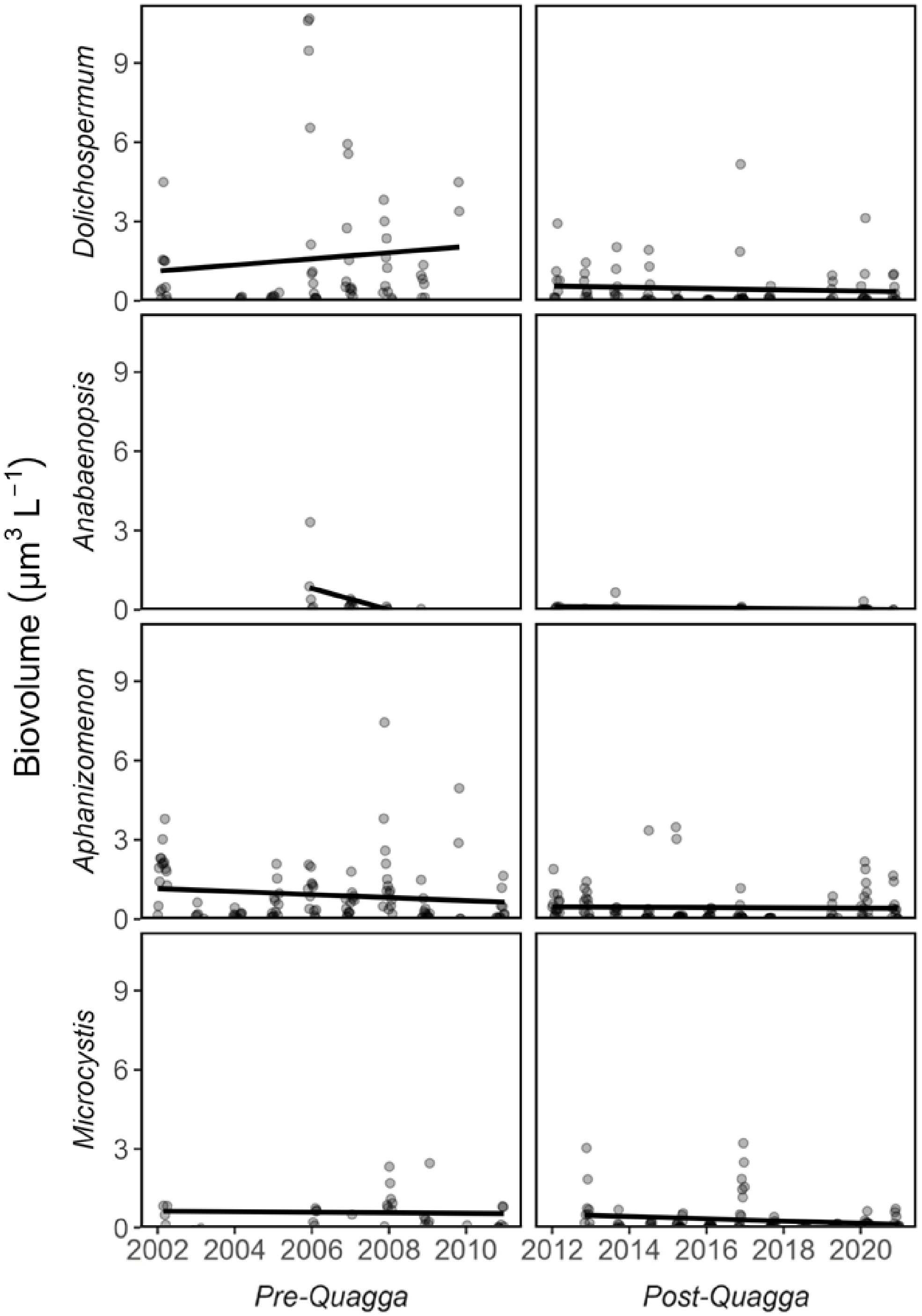


Supplementary Figure S5: Summer (July -September) cyanobacterial biovolume for the four most abundant species (*DolichospermumAnabaenopsis*, *Aphanizomenon* and *Microcystis*) in Lake Müggelsee before (2002-2011, left) and after (2012-2021) quagga mussel invasion.

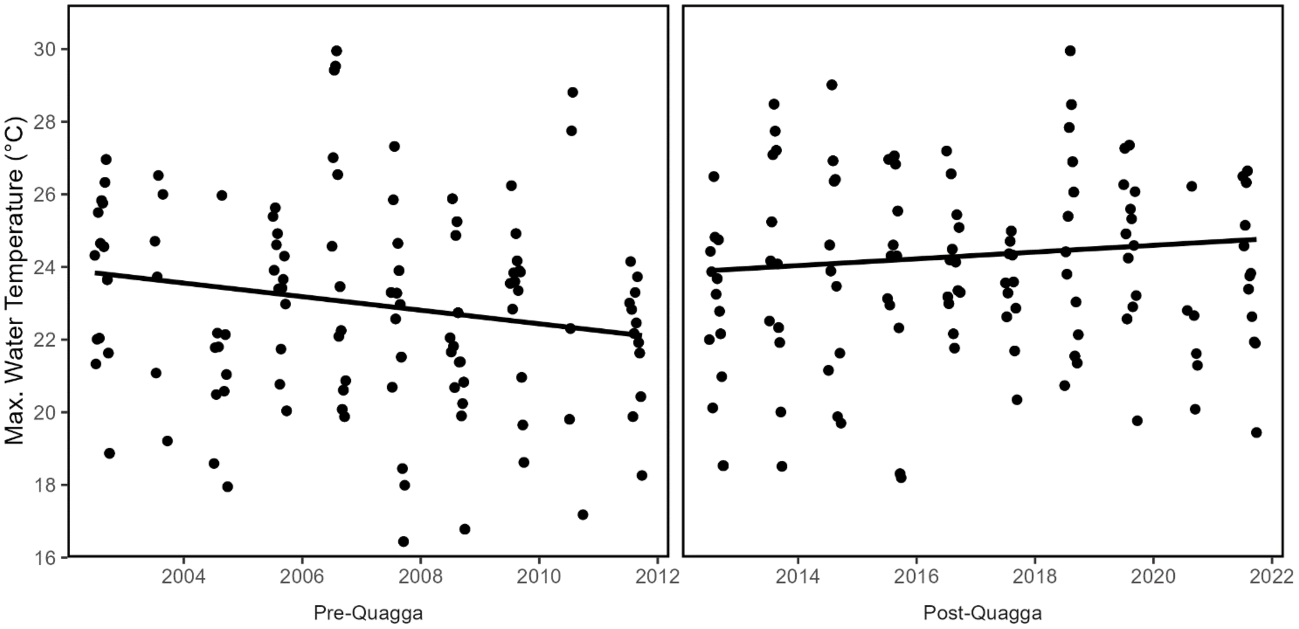


Supplementary Figure S6: Summer (July -September) maximum (Max.) water temperatures in the week before phytoplankton samplings in Lake Müggelsee (see Figure 3) before (2002-2011) and after (2012-2021) quagga mussel invasion.

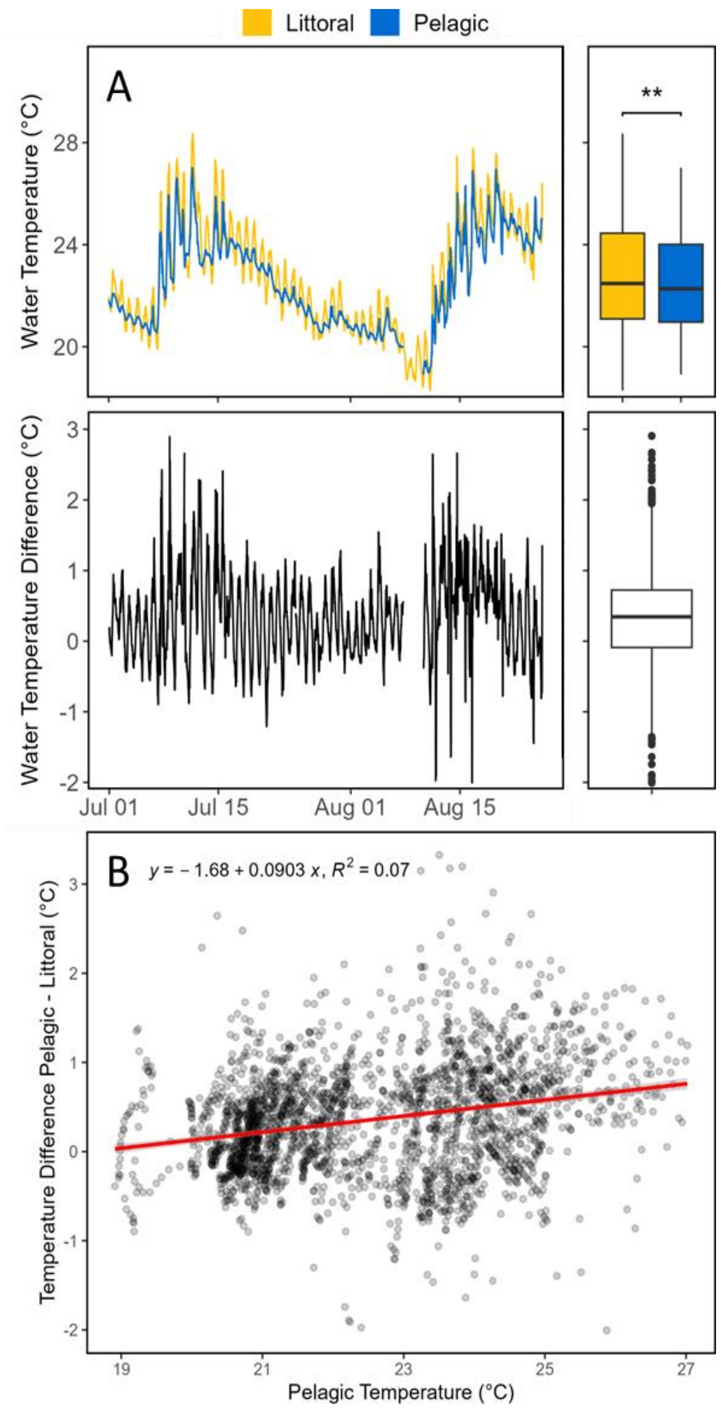


Supplementary Figure S7: Water temperature at littoral (yellow) and pelagic (blue) sites of Lake Müggelsee in 1 m water depth (t-test: $p < 0.01$) and differences between littoral and pelagic water temperatures (A) from July 01, 2023 until August 25, 2023. Water temperature difference along a gradient of pelagic water temperatures and linear regression (B).

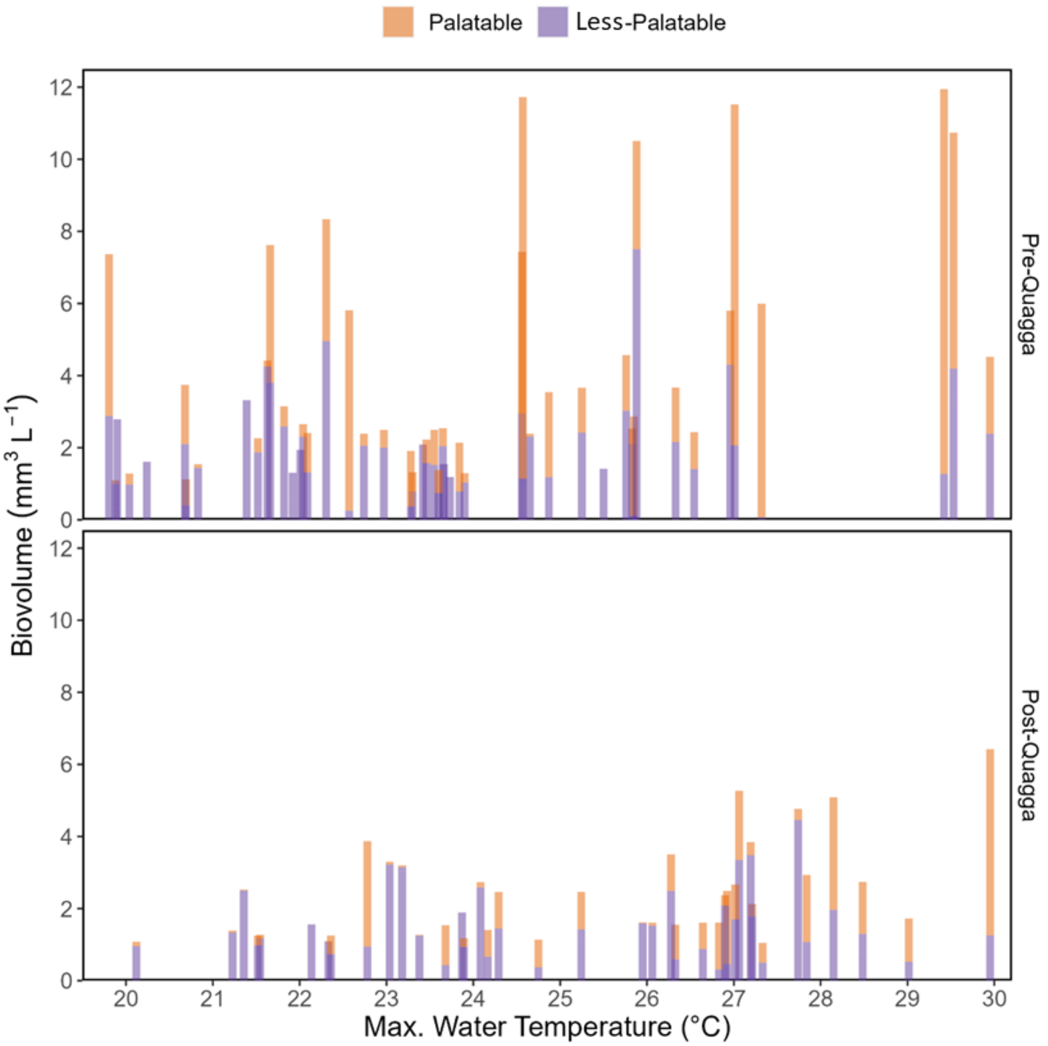


Supplementary Figure S8: Biovolumes of palatable (orange) and less-palatable (purple) cyanobacteria before (2002-2011, pre-quagga) and after (2012-2021, post-quagga) quagga mussel invasions in Lake Müggelsee plotted along a maximum (max.) water temperature gradient (measured during the week before phytoplankton sampling)

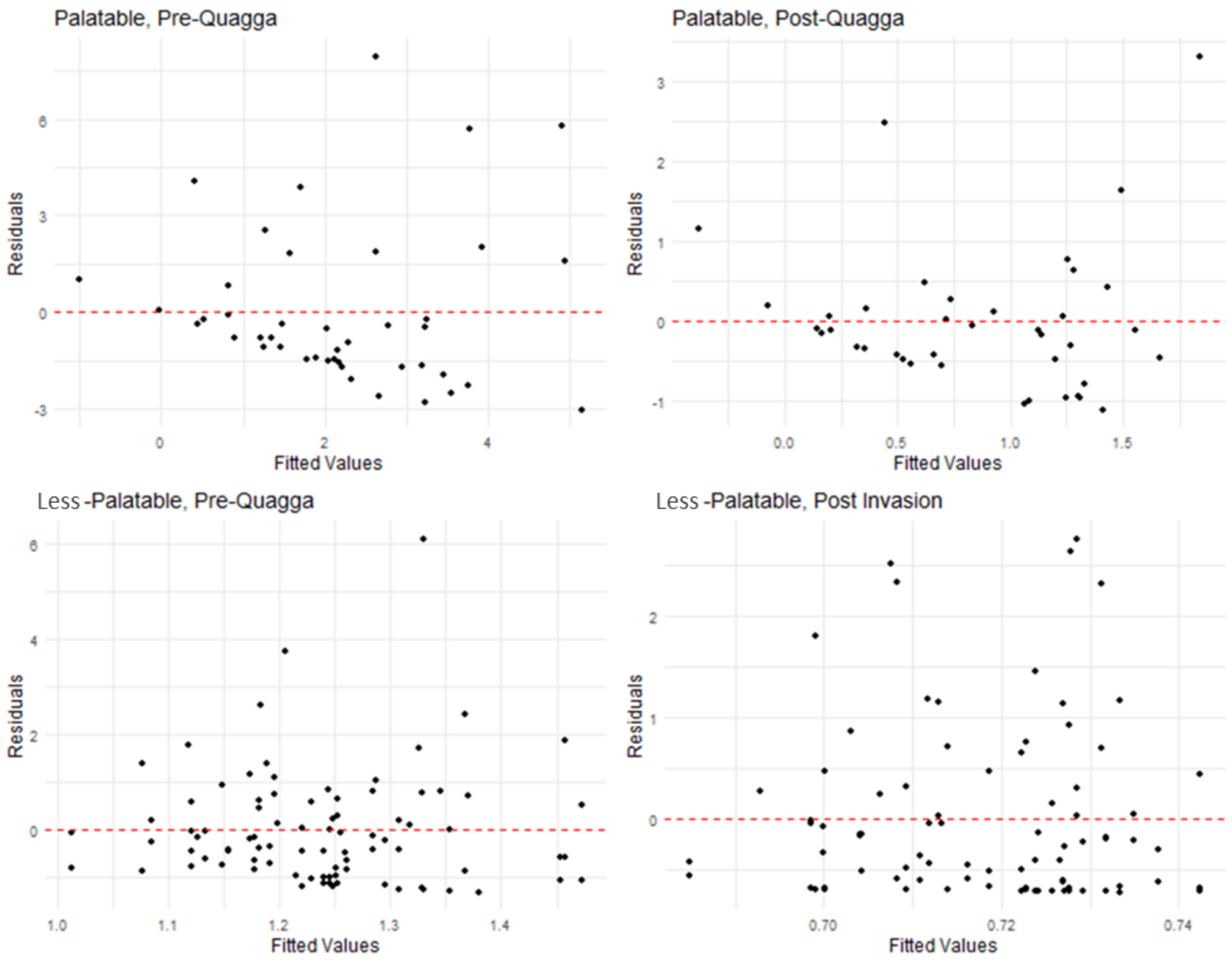


Supplementary Figure S9: Assessment of homoscedasticity in temperature-biovolume relationship cyanobacteria before (2002-2011, pre-quagga) and after (2012-2021, post-quagga) quagga mussel invasions in Lake Müggelsee (Figure 6). Residual plot showing the distribution of model residuals against fitted values from the linear regression of cyanobacterial biovolume on maximum water temperature. The x-axis represents the fitted (predicted) biovolume values from the model, while the y-axis shows the corresponding residuals (observed minus predicted values). The red dashed horizontal line at y = 0 represents perfect prediction.

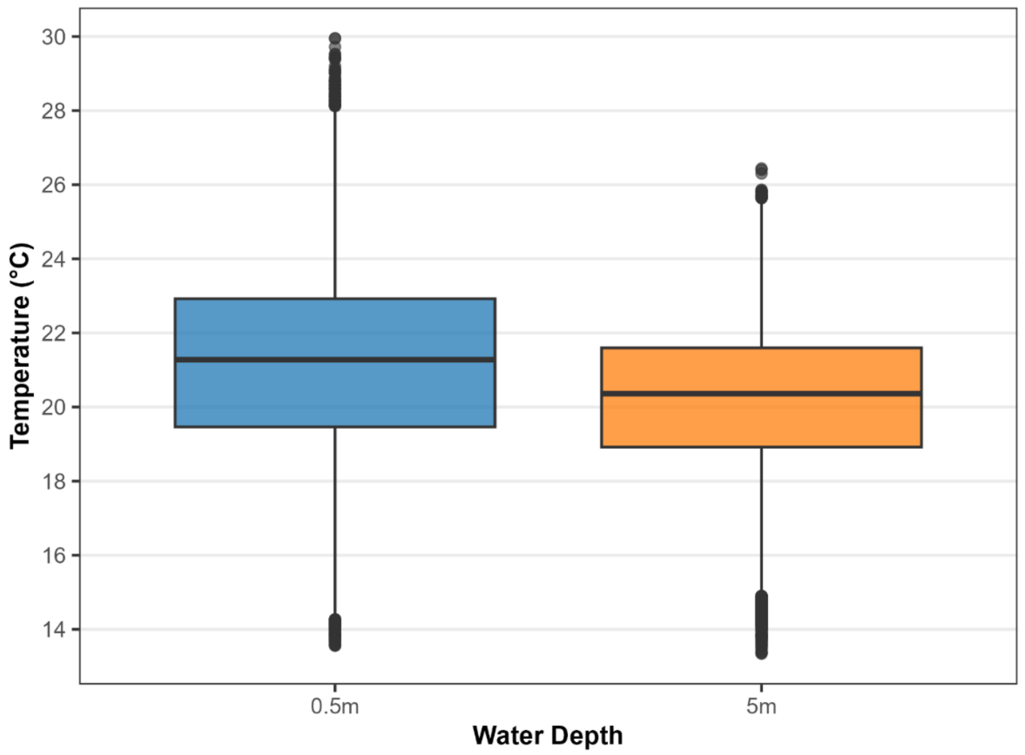


Supplementary Figure S10: Comparison of summer (Jul.-Sep.) water temperatures at 0.5 and 5 m depth in Lake Müggelsee.

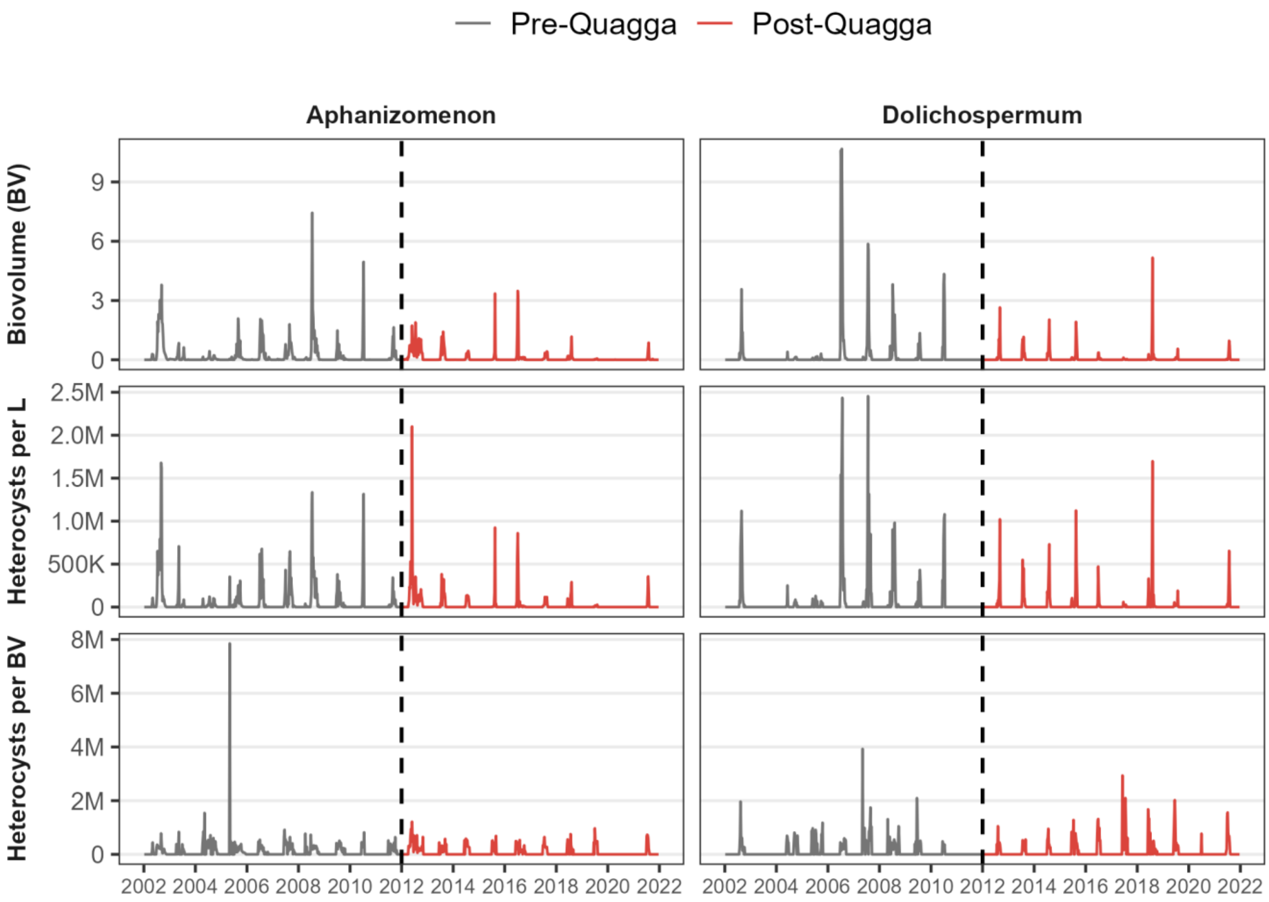


Supplementary Figure S11: Biovolumes, heterocyst abundance (per liter), and heterocyst frequency per biovolume (BV) of *Dolichospermum* (left column) and *Aphanizomenon* (right column) before (2002-2011, Pre-Quagga) and after (2012-2021, Post-Quagga) quagga mussel invasion in Lake Müggelsee. Black dashed line indicates quagga mussel invasion.

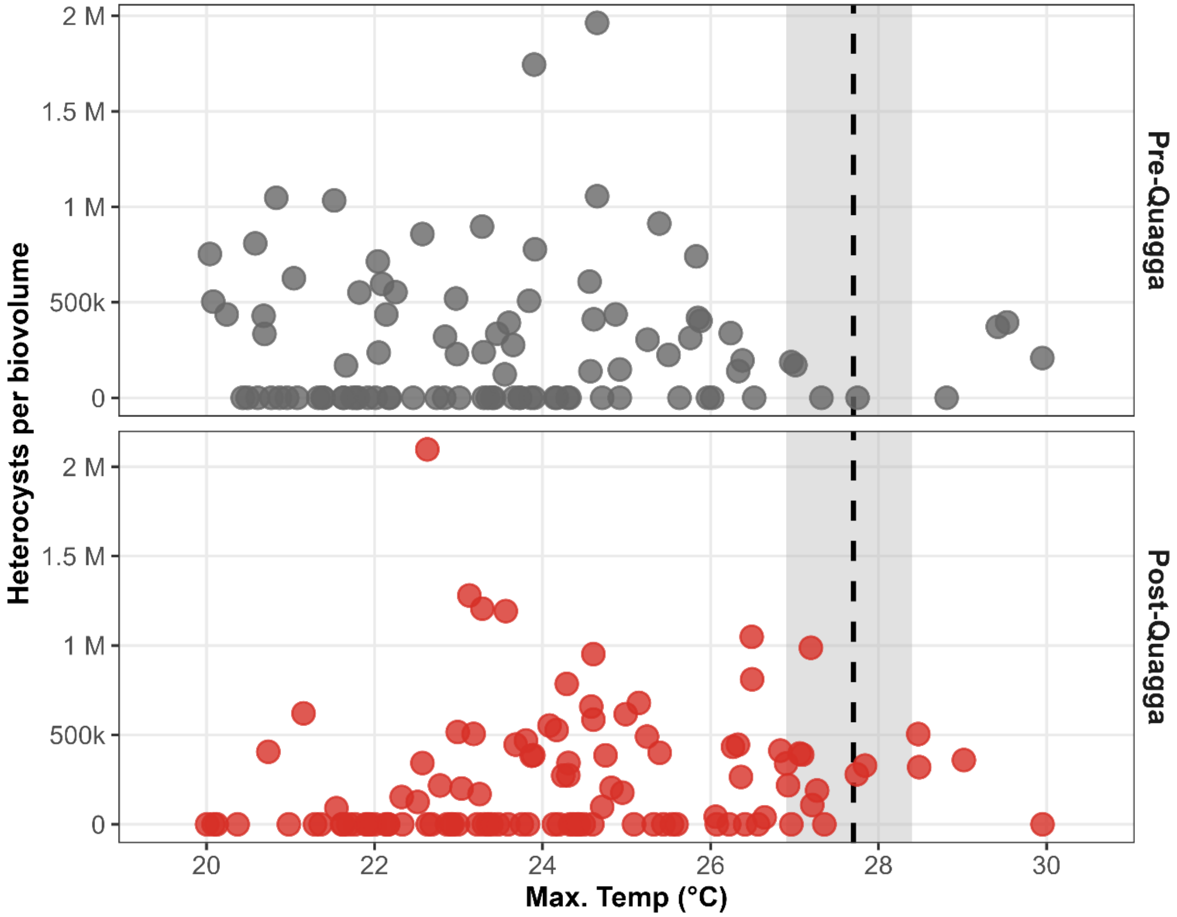


Supplementary Figure S12: Relationship between heterocyst frequency per biovolume (BV) of *Dolichospermum* and maximum (Max.) temperature of the previous week in Lake Müggelsee before (2002–2011, Pre-Quagga, top) and after (2012–2021, Post-Quagga, bottom) quagga mussel invasion. Points show weekly data; dashed black line indicates break-point and grey are the confidence interval from long-term data analysis in Figure 6.
